# Regulation of layered T cell tolerance mechanisms by the NR4A family is essential to preserve immune homeostasis and suppress autoimmunity

**DOI:** 10.1101/2021.04.28.441904

**Authors:** Ryosuke Hiwa, Hailyn V. Nielsen, James L. Mueller, Julie Zikherman

**Affiliations:** Division of Rheumatology, Rosalind Russell and Ephraim P. Engleman Arthritis Research Center, Department of Medicine, University of California, San Francisco, CA, 94143

**Keywords:** NUR77, NOR-1, *Nr4a1*, *Nr4a3*, Treg, T cell tolerance, anergy, negative selection

## Abstract

The NR4A family of orphan nuclear receptors (*Nr4a1-3*) plays redundant roles upstream of *Foxp3* to establish and maintain Treg identity; deletion of multiple family members in the thymus results in Treg deficiency and a severe inflammatory disease. Consequently, it has been challenging to isolate the functions of this family in other immune cells. Here we take advantage of a competitive bone marrow chimera strategy, coupled with conditional genetic tools, to rescue Treg homeostasis and unmask such functions. Unexpectedly, chimeras harboring *Nr4a1^−/−^ Nr4a3^−/−^* (DKO) bone marrow develop autoantibodies and a systemic inflammatory disease despite a replete Treg compartment of largely wild-type origin. This disease differs qualitatively from that seen with Treg-deficiency and is B cell-extrinsic. Negative selection of DKO thymocytes is profoundly impaired in a cell-intrinsic manner. Consistent with escape of self-reactive T cells into the periphery, DKO T cells with functional and phenotypic features of anergy accumulate in chimeric mice. Despite this, DKO T cells exhibit enhanced IL-2 production, implying a cell-intrinsic role for the NR4A family in peripheral T cell tolerance. These studies reveal roles for the NR4A family in multiple layered T cell tolerance mechanisms and demonstrate that each is essential to preserve immune homeostasis.

## INTRODUCTION

Since the initial discovery of regulatory T cells (Treg) (1, 2) and their formal recognition as a distinct T cell lineage dependent upon the transcription factor FOXP3 (3–5), it has been appreciated that they are absolutely essential for immune homeostasis and tolerance to self. Indeed, *Foxp3*-deficient mice and mice with a loss-of-function mutation in *Foxp3* (Scurfy) rapidly develop a frank autoimmune, lymphoproliferative, and myeloproliferative disease characterized by cytokine storm, immune cell infiltration, autoantibody production, and death typically by 4 weeks of age (6–9). Conversely, re-introducing Treg is sufficient to prevent this disease (6) and analogous immune therapies have been pioneered in humans (10). However, extensive *cell-intrinsic* mechanisms that operate in other immune cell lineages are also essential to maintain tolerance to self, including processes such as deletion, receptor editing (in the case of B cells), and hypo-responsiveness of self-reactive lymphocytes (termed anergy) (11–13).

Prior work has implicated a small family of orphan nuclear hormone receptors (encoded by *Nr4a1-3*) in several of these processes. Most notably, perhaps, NR4A family members play redundant roles upstream of *Foxp3* to maintain Treg identity and function; deletion of multiple family members in the thymus results in profound Treg deficiency and a severe “Scurfy-like” disease that phenocopies *Foxp3*-deficient mice (14, 15). As a result, it has been difficult to isolate redundant functions of this family in other immune cell populations. Yet this remains an important area to explore since the NR4A family are widely expressed and thought to be druggable targets that may facilitate manipulation of immune cell function in the context of autoimmune disease (16), tumor immunotherapy (17), and hematologic malignancies (18); indeed, such efforts are already underway despite these limitations. It is, therefore, critical to circumvent Treg dysfunction in order to isolate and define additional cell-intrinsic contributions of the NR4A family to immune cell homeostasis and tolerance.

*Nr4a1-3* (encoding NUR77, NURR1, and NOR-1, respectively) are rapidly upregulated in response to mitogenic stimuli, including antigen receptor ligation, and are thought to function as constitutively active transcription factors without a clear endogenous ligand (19). As a result, not only are these family members upregulated in T and B cells after acute antigen encounter, but also in Treg in the steady-state, in thymocytes undergoing negative selection, and in self-reactive, anergic, or exhausted lymphocytes in response to chronic antigen stimulation (14, 16, 17, 20–26). Indeed, the NR4A family has been argued to play a tolerogenic role in all these contexts. The NR4A family selectively restrains the survival and expansion of B cells that encounter antigen (signal 1) in the absence of co-stimulation (signal 2) (22, 27). Similarly, overexpression of either *Nr4a1* or *Nr4a3* mediates antigen-induced cell death in the thymus, while a dominant-negative transgenic (Tg) construct had the opposite effect (20, 28, 29). However, *Nr4a1^−/−^* mice exhibit extremely subtle defects in thymic negative selection (30, 31), suggesting possible redundancy among the family members. *Nr4a1* and *Nr4a3* also play non-redundant roles in peripheral conventional T cells (Tconv): most notable among these are roles for *Nr4a1* in CD4^+^ T cell anergy (24) and an additive role for all 3 family members in CD8^+^ T cell exhaustion (17). Finally, it has been argued that *Nr4a1* and *Nr4a3* redundantly maintain myeloid homeostasis since, in their absence, a myeloproliferative disease is observed (32). However, unmasking redundancy between NR4A family members in many of these contexts has been hampered by profound immune dysregulation that develops in the absence of functional Treg.

We sought to bypass this obstacle by generating competitive bone marrow (BM) chimeras harboring both wild-type (WT) cells (that could reconstitute a functional Treg compartment) and DKO cells (lacking both *Nr4a1* and *Nr4a3*) in order to isolate cell-intrinsic immune functions for the NR4A family. Unexpectedly, mixed chimeras harboring both WT and DKO BM rapidly developed anti-nuclear autoantibodies (ANA) and a systemic inflammatory disease, despite a replete Treg compartment of largely WT origin. The disease that developed in BM chimeras was B cell-extrinsic and qualitatively different from that in germline DKO mice. We found that negative selection of DKO thymocytes in competitive chimeras was profoundly impaired in a cell-autonomous manner. DKO Tconv cells with phenotypic and functional features of antigen experience and anergy accumulate in these chimeras, suggesting escape of self-reactive T cells into the periphery. However, self-reactive DKO CD4^+^ Tconv cells nevertheless exhibit exaggerated IL-2 production, suggesting that anergy is defective. Our findings unmask essential, redundant roles for the NR4A family in both central and peripheral T cell tolerance to maintain immune homeostasis.

## RESULTS

### Systemic immune dysregulation in mice with germline deficiency of *Nr4a1* and *Nr4a3*

*Nr4a1, 2*, and *3* are all expressed in thymocytes, Treg, and peripheral T cells, but the expression of *Nr4a2* is minimal under steady-state conditions (**Supplementary Figure 1A**; immgen.org). In order to unmask redundant functions of the NR4A family, we generated mice lacking germline expression of both *Nr4a1* and *Nr4a3*. We used *Nr4a1^fl/fl^* mice to generate *Nr4a1*-deficient mice with germline excision of the loxp-flanked locus and bred this with a CRISPR-generated *Nr4a3^−/−^* line that we recently described (27). An independently-generated line of *Nr4a1^−/−^* mice in widespread use has been reported to express a truncated NUR77 protein encoded by exon 2 of *Nr4a1* (30, 33). Our newly generated *Nr4a1^−/−^ Nr4a3^−/−^* mice (germline DKO, denoted as gDKO here-in) do not express exon 2 of *Nr4a1* consistent with the prior analysis of *Nr4a1^fl/fl^* mice (33).

gDKO mice were born at Mendelian ratios but exhibited severe runting (**Figure 1A**) and invariably died before 4 weeks of age, consistent with observed mortality in an independent gDKO line generated with distinct *Nr4a1* and *Nr4a3* mutant alleles (32). As previously reported for CD4-cre *Nr4a^fl/fl^ Nr4a3^−/−^* mice (14), our new line of gDKO mice exhibit near-complete loss of FOXP3^+^ Treg in both thymus and periphery (**Figures 1B-D, Supplementary Figure 1B**). Concurrently, we observed expansion of a unique population of CD4^+^CD25^+^ FOXP3^−^ T cells in the periphery that may represent T cells that have either lost or failed to upregulate expression of FOXP3 described elsewhere (**Figures 1B, E, Supplementary Figures 1B-D**) (14, 15, 34). Importantly, and consistent with prior reports, neither *Nr4a1^−/−^* nor *Nr4a3^−/−^* single knockout (SKO) mice exhibit Treg loss, expansion of this unique cell population, or frank disease (**Figures 1B-E, Supplementary Figures 1B-D**) (14).

**Figure 1.**
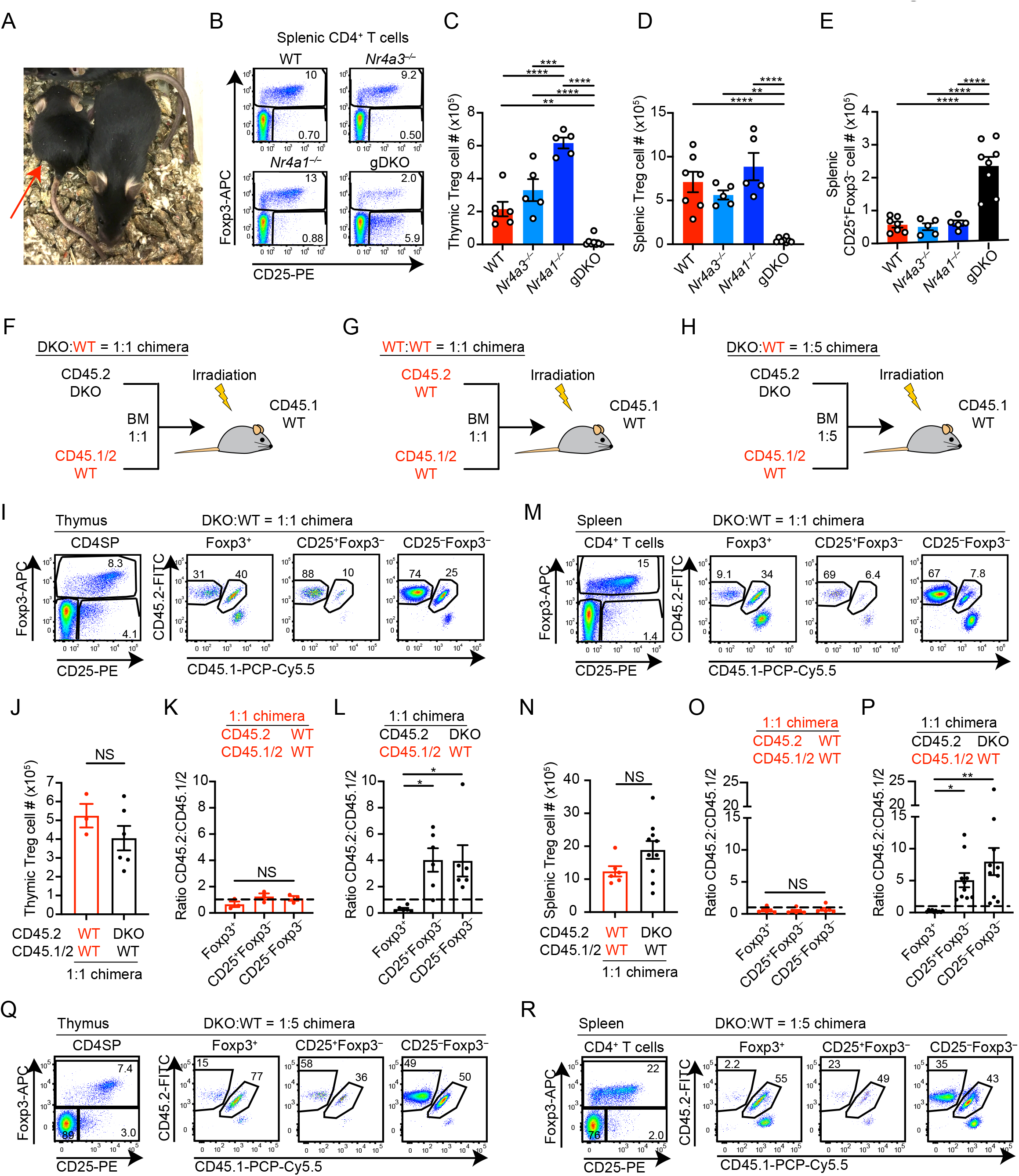
Systemic immune dysregulation and Treg-defect in mice with germline deficiency of *Nr4a1* and *Nr4a3*. A. *Nr4a1^−/−^Nr4a3^−/−^* (gDKO) mouse (red arrow) is small with irritated tail skin compared to healthy littermate, 4 w; Representative of n = 8. B. Representative flow plots show splenic CD4^+^ T cells with Treg gate (FOXP3^+^) in mice of each genotype as shown. Plots are representative of n = 5 mice/genotype. C-E. Quantification of thymic Treg (C), splenic Treg (D) and CD4^+^CD25^+^FOXP3^−^ (E) cell number as gated in B (n ≧ 5 biological replicates, 3 to 4-week-old gDKO and 5 to 6-week-old mice with the other genotypes were analyzed). F-H. Schematics depict competitive bone marrow chimera design. CD45.2 DKO (F, H) or CD45.2 WT (G) bone marrow cells were mixed with CD45.1/2 WT bone marrow cells in 1:1 (F, G) or 1:5 (H) ratio, and transferred into lethally irradiated CD45.1 WT hosts. I, M. Representative flow plots show thymic CD4SP (I) or splenic CD4^+^ T cell populations in 1:1 DKO:WT chimeras to identify genetic composition of indicated subpopulations. Plots are representative of 6 (I) or 10 (M) chimeras. J-P. Quantification of thymic (J) or splenic (N) Treg cell number in 1:1 chimeras. Ratio of CD45.2 to CD45.1/2 for thymic (K, L) or splenic (O, P) Treg, CD25^+^FOXP3^−^ and CD25^−^FOXP3^−^ cells in 1:1 WT:WT (K, O) and DKO:WT chimera (L, P). Ratios were normalized to DP thymocytes. Data were from 6–10 w post-transplant chimeras (n ≧ 3 (J-L) or 6 (N-P). Q, R. Representative flow plots show thymic CD4SP (Q) or splenic CD4^+^ T cell (R) population in 1:5 DKO:WT chimera. Plots are representative of 3 chimeras (quantified in Supp. Fig 1F, G). Graphs depict mean +/- SEM. Statistical significance was assessed by one-way ANOVA with Tukey’s test (C-E, K, L, O, P) or two-tailed unpaired Student’s t-test (J, N). *P < 0.05; **P < 0.01; ***P < 0.001; ****P < 0.0001. NS, not significant.

### Cell intrinsic Treg defect in the absence of *Nr4a1* and *Nr4a3*

Systemic inflammatory disease and associated profound thymic atrophy preclude the study of thymic development and mature CD4^+^ Tconv cells in gDKO mice (**Supplementary Figure 1E**). Similar mortality and Treg deficiency observed in both germline (32) and CD4-cre conditional mouse lines (14) suggested to us that disease might be due to Treg deficiency in both settings. We reasoned that restoring functional Treg could unmask cell-intrinsic functions of NR4A family in other immune cell populations. To do so, we next generated competitive chimeras to allow WT donor BM to reconstitute a functional Treg compartment. Equal proportions of congenically marked donor BM from CD45.2 gDKO and CD45.1/2 WT mice was transplanted into lethally irradiated CD45.1 BoyJ recipients (**Figure 1F**). In parallel, we generated control chimeras in which CD45.1 hosts were reconstituted with a mixture of CD45.2 WT and CD45.1/2 WT BM (**Figure 1G**). In addition, we also generated mixed chimeras with a low proportion of gDKO donor BM (1:5 ratio) in order to further ensure development of largely WT Treg compartment (**Figure 1H**). We assessed reconstitution and immune phenotypes of chimeras at sequential time points between 6-14 weeks after transplant.

Consistent with studies of CD4-cre chimeras, we observed a profound cell-intrinsic disadvantage for DKO Treg in the thymus (**Figures 1I-L**) and spleen (**Figures 1M-P**) when compared to CD4SP thymocytes (14). This was not attributable to CD45 allotype since no disadvantage for CD45.2 thymocytes was seen in WT:WT chimera (**Figures 1K, O**). Conversely, we observed enrichment of DKO cells in CD4^+^CD25^+^ FOXP3^−^ compartment (**Figures 1I, L, M, P**). Similar results were reproduced with DKO:WT 1:5 chimera (**Figure 1Q, R, Supplementary Figures 1F, G**). FOXP3 and CD25 expression in DKO Treg was reduced (**Supplementary Figures 1H-K**), consistent with a role for the NR4A family in ‘maintenance’ of Treg identity (25). Most importantly, the Treg compartment was restored and the number of Treg was comparable between DKO:WT chimera and WT:WT chimera (**Figures 1J, N**). We also confirmed Treg were largely reconstituted from WT donor in DKO:WT 1:5 chimera (**Figures 1Q, R**). This allowed us to explore the cell-intrinsic roles of the NR4A family in other immune cell types.

### Thymic atrophy is rescued in competitive chimeras

gDKO mice exhibit severe thymic atrophy with marked reduction of all thymocyte subsets and disproportionate loss of DP thymocytes (**Figures 2A, B**). We postulated that this was an indirect consequence of Treg deficiency and systemic inflammation in gDKO mice, and reasoned that thymic development could be assessed in competitive chimeras. Indeed, profound thymic atrophy is partially rescued in DKO:WT 1:1 chimeras within the first 6 weeks of reconstitution (**Supplementary Figure 2A**). However, progressive thymic atrophy was observed over time in DKO:WT 1:1 chimeras (relative to WT:WT control chimera) (**Supplementary Figure 2A**). This suggested to us that thymic atrophy was incompletely repaired by WT Treg and led us to focus on early time points to isolate cell-intrinsic roles for the NR4A family during thymic development (6 weeks post-transplant). In addition, we assessed competitive chimeras harboring 1:5 ratio of DKO:WT donor BM in which both Treg number and thymic cellularity were completely rescued at 6 weeks post-transplant (**Supplementary Figures 2A-C**).

**Figure 2.**
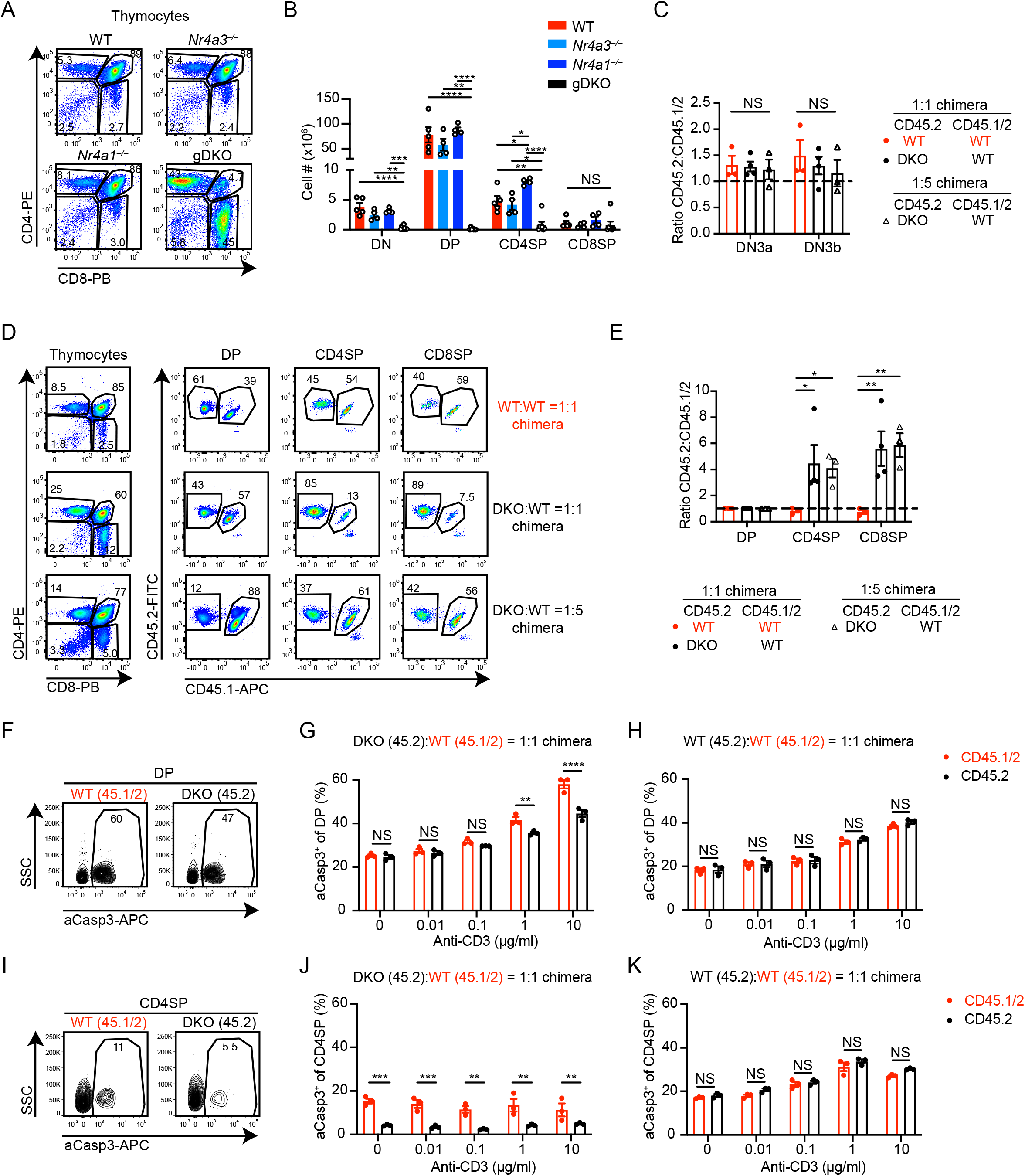
DKO thymocytes have a cell-intrinsic defect in negative selection. A. Representative flow plots show thymic subsets in WT, *Nr4a3^−/−^*, *Nr4a1^−/−^* and gDKO mice. Plots are representative of ≧ 4 mice/genotype. B. Quantification of thymic subset cell number as gated in (A); (n ≧ 4 biological replicates, 3 to 4-week-old gDKO and 5 to 6-week-old mice with the other genotypes were analyzed). C. Ratio of CD45.2 to CD45.1/2 thymocytes among thymic DN3a and DN3b subsets from chimeras (as gated in Supp. Fig. 2D). Ratios normalized to DN2 subset (n = 3-4 biological replicates). D. Representative flow plots show thymic subsets in competitive chimera. Plots are representative of ≧ 3 mice/genotype. E. Ratio of CD45.2 to CD45.1/2 thymic subsets as gated in (D) in chimeras normalized to DP thymocytes (n ≧ 3 biological replicates). Data in C-E were from 6–7 w post-transplant chimeras. F-K. Thymocytes from 1:1 DKO:WT chimeras were cultured with varying doses of plate-bound anti-CD3 and 2 μg/ml of anti-CD28 for 24 h. Cells were stained to detect CD4/CD8 surface markers, followed by permeabilization and detection of active Caspase3 (aCasp3) via flow cytometry. Representative plots show aCasp3 expression in WT CD45.1/2 and DKO CD45.2 DP (F) and CD4SP (I) thymocytes from 1:1 DKO chimeras cultured with 10 μg/ml anti-CD3. Quantification of frequency of aCasp3^+^ cells among DP (G, H) or CD4SP (J, K) in 1:1 DKO:WT (G, J) or WT:WT (H, K) chimeras cultured as described above (n = 3 biological replicates). Graphs depict mean +/- SEM. Statistical significance was assessed by one-way (B) or two-way (C, E) ANOVA with Tukey’s test or two-tailed unpaired Student’s t-test with the Holm–Šídák method (G, H, J, K). *P < 0.05; **P < 0.01; ***P < 0.001; ****P < 0.0001. NS, not significant.

### NR4A expression is dispensable for thymic β-selection

We previously showed, using a fluorescent reporter of *Nr4a1* transcription (NUR77-eGFP), that GFP is upregulated at the β-selection checkpoint during thymic development, raising the possibility that *Nr4a1* and family members might play a functional role here (21). Immature double negative (DN) thymocytes (lacking both CD4 and CD8 expression) first undergo recombination of the TCRβ chain, which then pairs with the pre-TCRα chain to signal in an antigen (Ag)-independent manner at the ‘β-selection’ checkpoint (35). This occurs during the DN3 stage of development; pre-selection DN3a thymocytes are defined as CD25^hi^CD44^lo^ forward scatter (FSC)-low thymocytes, while DN3b thymocytes that have traversed this checkpoint successfully express the same surface markers but are larger (FSC-high) (36) (**Supplementary Figure 2D**). We probed β-selection in both 1:1 and 1:5 DKO:WT competitive chimeras, but identified no advantage for either DKO or WT CD45.2 cells relative to competitor CD45.1/2 WT cells (**Figure 2C**).

### DKO thymocytes have a profound cell-intrinsic defect in negative selection

Studies of two independent NUR77-eGFP reporter lines as well as transcriptional analysis have shown that *Nr4a* genes are upregulated at the positive selection checkpoint and are especially enriched among thymocytes undergoing negative selection (21, 23, 25, 37). Although overexpression of full-length and truncated dominant-negative constructs suggested that NUR77 and NOR-1 redundantly mediate thymic negative selection (20, 29, 38, 39), it has not been possible to date to study negative selection in animals lacking both family members because profound immune dysregulation leads to severe thymic atrophy (**Figure 2B**, **Supplementary Figure 1E**). Indeed, thymic negative selection is largely intact in *Nr4a1^−/−^* mice with only subtle defects, consistent with putative redundancy among family members (30, 31).

We reasoned that DKO:WT competitive chimeras could unmask cell-intrinsic, redundant functions of the NR4A family during thymic selection. Indeed, we observe a striking advantage for DKO cells in CD4SP and CD8SP subsets relative to DP in both 1:1 and 1:5 chimeras (**Figures 2D, E**), suggesting either enhanced positive selection or impaired negative selection. However, we do not see an advantage for DKO cells in post-selection DP thymocytes relative to pre-selection DP thymocytes, arguing against a role during positive selection (**Supplementary Figures 2E, F**).

Studies of *in vitro* and Tg model systems suggested a role for the NR4A family in TCR-induced cell death (20, 28, 29, 38, 39). Therefore, we hypothesized that the striking advantage for DKO thymocytes during development (**Figures 2D, E**) might be due to escape from negative selection. To further test this hypothesis, we assessed Ag-induced apoptosis by detection of activated Caspase 3 (aCasp3) expression in thymocytes from chimeras. We observed reduced aCasp3^+^ DKO relative to WT thymocytes after *in vitro* TCR-stimulation (**Figures 2F, G, I, J**). By contrast, we saw no difference between donors in control WT:WT chimeras (**Figures 2H, K**). Notably, we also saw no significant difference in aCasp3 expression in SKO thymocytes relative to co-cultured WT (**Supplementary Figures 2G, H**). Moreover, mixed chimeras generated with either *Nr4a1^−/−^* or *Nr4a3^−/−^* SKO mice revealed only a small competitive advantage for CD8SP cells, suggesting a largely redundant role for these family members during negative selection that is only unmasked when both family members are lost (40) (**Supplementary Figure 2I**). We conclude that DKO thymocytes escape negative selection, and show for the first time that this is a profound effect in a physiological context, independent of either a TCR Tg or NR4A mis-expression.

### Myeloproliferative disorder in DKO mice is cell-extrinsic

Previous studies report that a severe myeloproliferative disorder develops in the first weeks of life in independently generated gDKO mice (32). This was not seen in SKO animals lacking only one *Nr4a* family member, although mice lacking three out of four *Nr4a* alleles (i.e. *Nr4a1^+/–^Nr4a3^−/−^* or *Nr4a1^−/−^Nr4a3^+/–^*) did eventually succumb to a similar disease at much later time points (41). Consistent with this, we observed profound expansion of side-scatter (SSC)-high cells infiltrating all hematopoietic tissues and lymphoid organs in gDKO mice; this included not only BM and spleen (**Supplementary Figure 3**), but was especially pronounced in lymph nodes and thymus (**Figures 3A-D**). These SSC^hi^ cells are CD11b^+^ but largely Gr1^−^. Since Treg-deficient animal models like Scurfy and *Foxp3*-deficient mice similarly exhibit myeloid expansion (6, 9, 42, 43), we hypothesized that the myeloproliferative disorder observed in gDKO animals was a non-cell-autonomous effect of Treg deficiency. Consistent with this possibility, myeloid expansion is observed in CD4-cre *Nr4a1^fl/fl^ Nr4a3^−/−^* mice, but not in mixed chimeras generated with WT donor BM (14). Resolving this question with gDKO cells has important implications since it has been argued that the NR4A family may represent important cell-intrinsic suppressors of myeloid leukemic diseases and drug development directed at this application is under way (18, 32, 41). Indeed, in our DKO:WT chimeras, myeloid expansion is suppressed (even after 12 weeks of reconstitution) and DKO cells exhibit no competitive advantage in these compartments (**Figures 3A-I, Supplementary Figure 3A, B, D, E**). We did observe a minor infiltration of SSC^hi^ CD11b^+^ cells into the thymus of 1:1 DKO:WT but not WT:WT chimeras, and here as well the effect of *Nr4a*-deficiency was cell-extrinsic (**Figures 3D, H, I**). Taken together, these data support our hypothesis that the myeloproliferative disorder observed in gDKO animals is due to a cell-extrinsic impact of *Nr4a* deletion in Treg.

**Figure 3.**
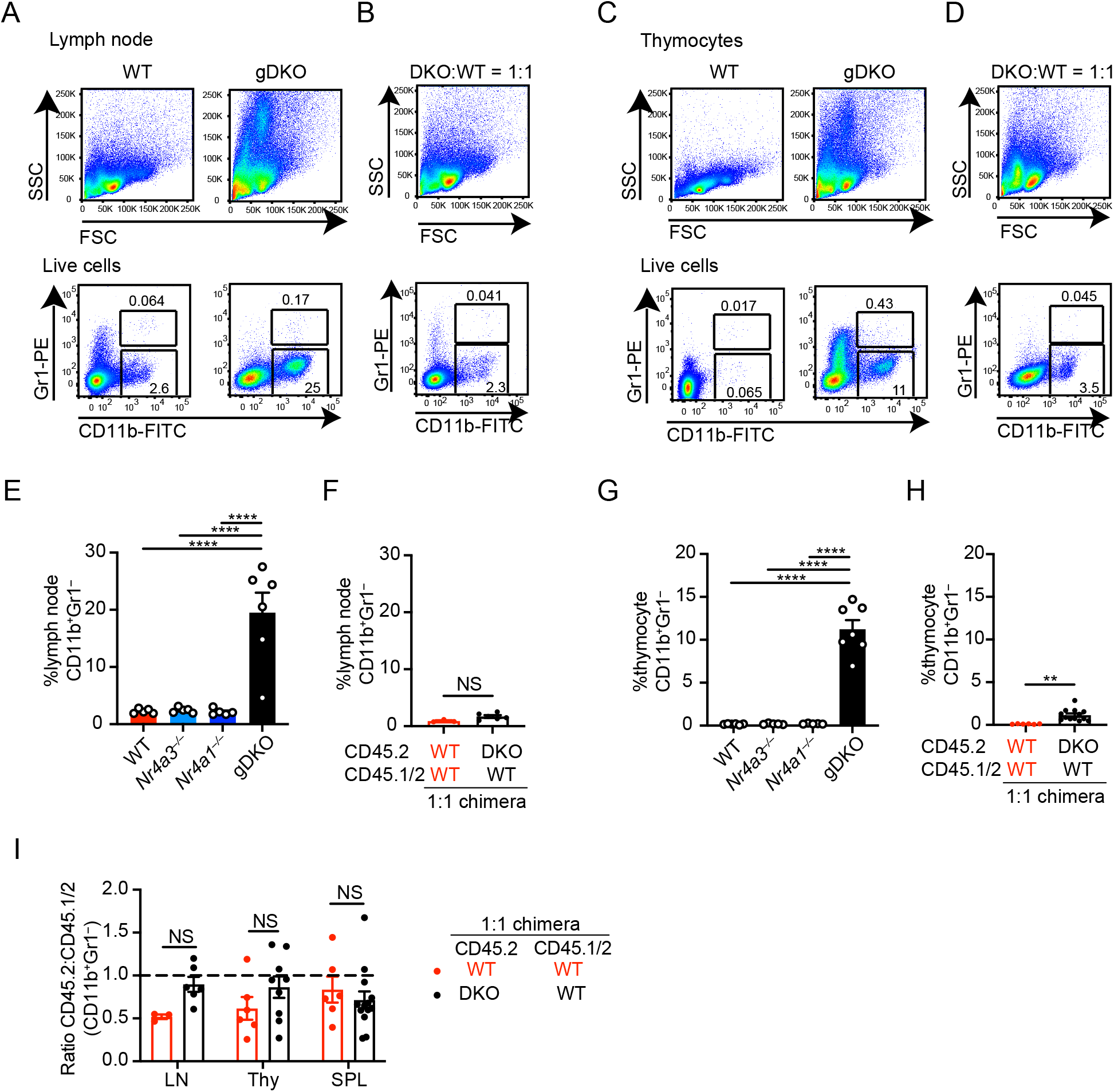
Myeloproliferative disorder in DKO mice is cell-extrinsic. A-D. Lymph nodes cells (A, B) and thymocytes (C, D) from WT and gDKO mice (A, C) or 1:1 DKO:WT chimeras (B, D) were stained to detect myeloid population determined by CD11b and Gr1 expression. Shown are representative plots of at least 5 mice. E, G. Quantification of CD11b^+^Gr1^−^ cells in lymph nodes (E) and thymocytes (G) from WT, *Nr4a3^−/−^*, *Nr4a1^−/−^* and gDKO mice (n ≧ 5 biological replicates, 3 to 4-week-old gDKO and 5 to 6-week-old mice with the other genotypes were analyzed). F, H. Quantification of CD11b^+^Gr1^−^ cell frequency in lymph nodes (F) and thymocytes (H) from WT:WT = 1:1 and DKO:WT = 1:1 chimera (n ≧ 3 biological replicates). L. Ratio of CD45.2 to CD45.1/2 for CD11b^+^Gr1^−^ cells in lymph nodes, thymus and spleen from WT:WT = 1:1 and DKO:WT = 1:1 chimera (n ≧ 3 biological replicates). Graphs depict mean +/- SEM. Statistical significance was assessed by one-way ANOVA with Tukey’s test (E, G), two-tailed unpaired Student’s t-test with (I) or without (F, H) the Holm–Šídák method. **P < 0.01; ****P < 0.0001. NS, not significant.

### Abnormal B cell homeostasis in DKO mice is cell-extrinsic

Like other Treg deficient models, gDKO mice exhibit spontaneous polyclonal B cell activation and differentiation under steady-state conditions (**Figures 4A, B**, **Supplementary Figures 4A-F**). We previously reported that *Nr4a1* expression scales with both acute and chronic antigen stimulation in B cells (21, 22, 27). More recently, we identified a cell-intrinsic role for the NR4A family in restraining Ag-induced B cell expansion when co-stimulatory signals are absent or limiting, including in the context of B cell tolerance (22, 27). We therefore sought to determine to what extent spontaneous B cell activation and differentiation in gDKO mice (under homeostatic conditions) were attributable to a B cell-intrinsic role for the NR4A family.

**Figure 4.**
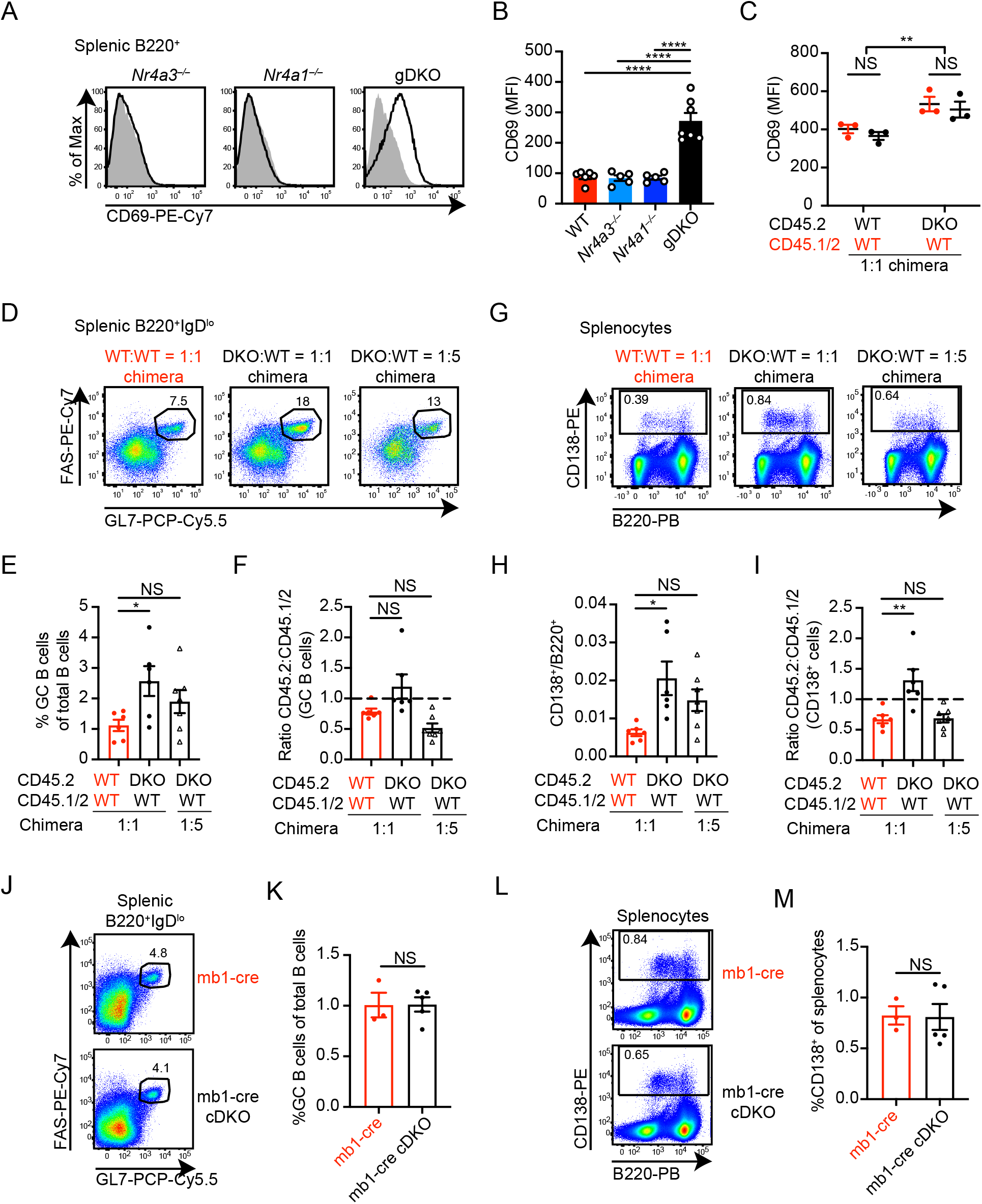
Abnormal B cell homeostasis in DKO mice is cell-extrinsic. A. Representative flow plots showing CD69 expression on splenic B cells from WT mice (shaded gray histogram) with B cell gate from *Nr4a3^−/−^*, *Nr4a1^−/−^* and gDKO mice overlaid. B. Quantification of CD69 MFI on splenic B cells from mice in (A) (Data in A, B represent n ≧ 5 biological replicates, 3 to 4-week-old gDKO and 5 to 6-week-old mice with the other genotypes were analyzed). C. Quantification of CD69 MFI on splenic B cells of each donor genotype in competitive chimeras (n = 3 biological replicates). D. Representative flow plots show FAS^hi^GL7^+^ GC B cells pre-gated on B220^+^IgD^lo^ splenocytes from competitive chimeras. E, F. Frequency of GC B cells among total B cells (E) and ratio of CD45.2 to CD45.1/2 GC B cells normalized to mature naïve B220^+^IgD^hi^ B cells in spleen from competitive chimeras as gated in D (data in D-F represent n ≧ 6 biological replicates). G. Representative flow plots show CD138^+^ gate among splenocytes from competitive chimeras. H, I. Ratio of CD138^+^ to B220^+^ cells and ratio of CD45.2 to CD45.1/2 CD138^+^ cells normalized to B220^+^CD138^−^ cells in spleen from competitive chimeras as gated in G (data in G-I represent n ≧ 6 biological replicates). J, L. Representative flow plots show GC B cells pre-gated on B220^+^IgD^lo^ (J) and CD138^+^ cells (L) in spleen from mb1-cre and mb1-cre *Nr4a1^fl/fl^ Nr4a3^−/−^* (cDKO) non-competitive chimeras 40 weeks after irradiation and transfer. K, M. Frequency of GC B cells among total B cells (K) and CD138^+^ cells among splenocytes (M) as described for J and L above (n ≧ 3 biological replicates). Graphs depict mean +/- SEM. Statistical significance was assessed by one-way ANOVA with Tukey’s test (B) or Dunnett’s test (E, F, H, I), or two-tailed unpaired Student’s t-test with (C) or without (K, M) the Holm–Šídák method. *P < 0.05; **P < 0.01; ****P < 0.0001. NS, not significant.

Consistent with our prior observations, DKO:WT chimeras did not reveal a competitive advantage or disadvantage for DKO cells during splenic B cell development apart from a subtle disadvantage in the MZ compartment (**Supplementary Figure 4G**) (27). B cells in 1:1 DKO:WT chimeras expressed higher levels of activation markers than B cells in WT:WT chimera (**Figure 4C, Supplementary Figures 4H, I**). However, this did not differ between donor populations within individual chimeras, suggesting a B cell-extrinsic effect. We observed expansion of GC B cells and CD138^+^ cells in DKO:WT chimeras relative to WT:WT control chimeras, but this was largely B cell-extrinsic (**Figures 4D-I**). Similarly, no expansion of the GC or CD138^+^ compartment was evident under steady-state conditions in mice lacking *Nr4a1/3* exclusively in the B cell compartment (mb1-cre *Nr4a1^fl/fl^ Nr4a3^−/−^*), even when aged to 40 weeks (**Figures 4J-M**). Nor could we detect an advantage for mb1-cre DKO B cells in a competitive setting (**Supplementary Figures 4J-L)**. We conclude that there is evidence of a spontaneous polyclonal B cell activation and differentiation in gDKO chimeras, but it is a largely B cell-extrinsic effect.

### Restoring the Treg compartment does not rescue DKO CD8 T cell homeostasis in mixed chimeras

Progressive thymic atrophy (**Supplementary Figure 2A**) and spontaneous B cell-extrinsic activation and differentiation in DKO:WT chimeras (**Figure 4**) suggested the development of a systemic autoimmune and inflammatory state despite replete and largely WT Treg compartment. We next probed the mature T cell compartment to understand the source of this immune dysregulation. gDKO mice exhibit expanded effector-memory compartment and nearly complete loss of naïve CD8^+^ T cells (**Figures 5A, B, Supplementary Figures 5A, B**). Indeed, the requirement for Treg to maintain CD8^+^ Tconv homeostasis by consuming IL-2, and CD4^+^ Tconv homeostasis through direct and cognate interactions, are among the most well-appreciated functions of Treg (44, 45).

**Figure 5.**
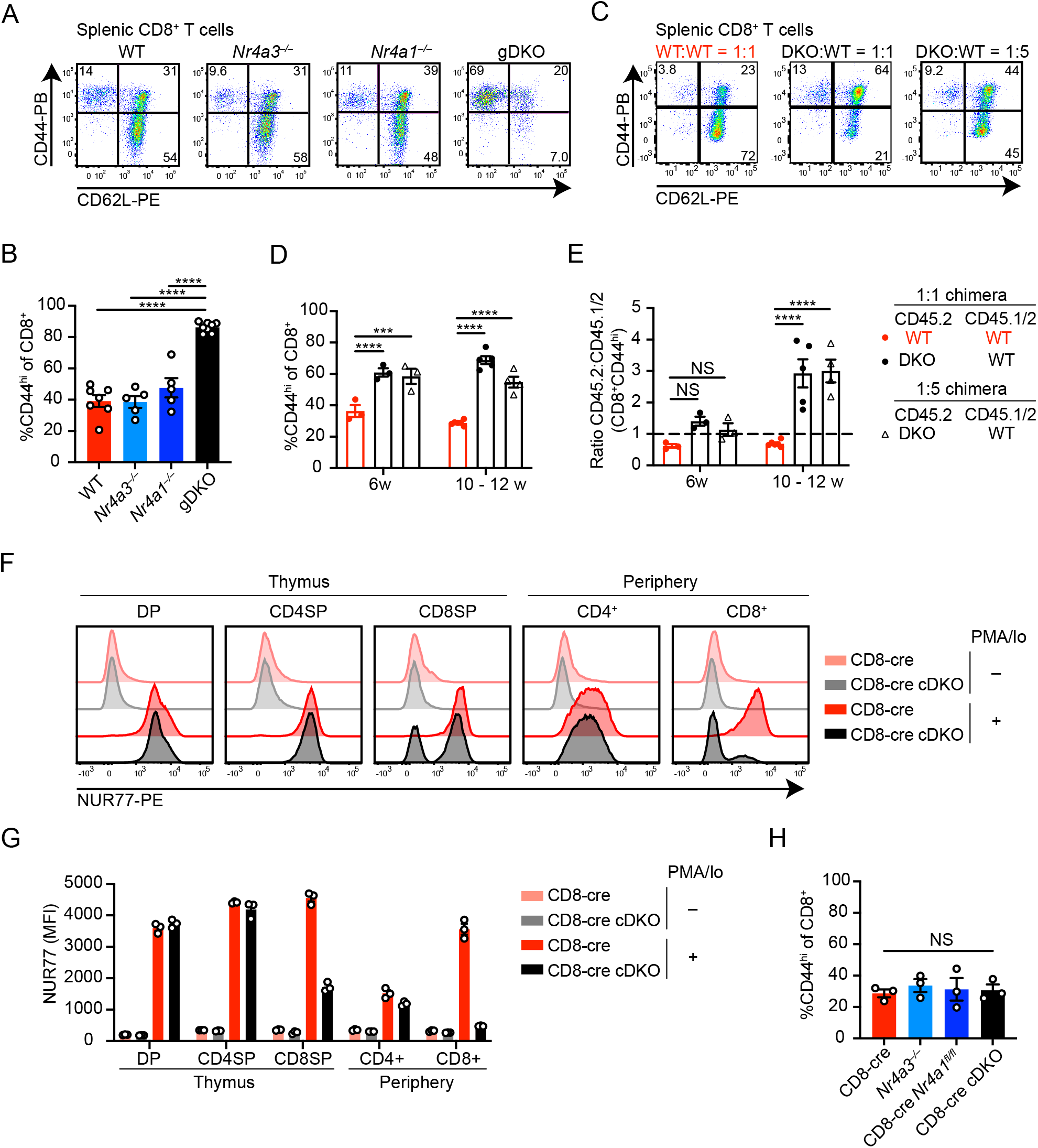
Reconstitution of Treg compartment does not restore CD8^+^ T cell homeostasis in DKO chimeras. A. Splenocytes from WT, *Nr4a3^−/−^*, *Nr4a1^−/−^* and gDKO mice were stained to detect CD8^+^ T cell subsets on the basis of CD44 and CD62L expression. Plots are representative of at least 5 mice. B. Quantification of frequency of splenic CD44^hi^ CD8^+^ T cells as gated in A (n ≧ 5 biological replicates, 3 to 4-week-old gDKO and 5 to 6-week-old mice with the other genotypes were analyzed). C. Representative flow plots showing the peripheral CD8^+^ T cell subsets in competitive chimeras, as described for A above. Plots are representative of at least 7 biological replicates. D. Quantification of frequency of splenic CD44^hi^ CD8^+^ T cells from competitive chimera as gated in C at varied time points following irradiation and BM transfer (n ≧ 3 biological replicates). E. Ratio of CD45.2 to CD45.1/2 for CD8^+^CD44^hi^ population gated in C, quantified in D. Ratio is normalized to naïve CD8^+^CD44^lo^CD62L^hi^ gate (n ≧ 3 biological replicates). F. Thymocytes and splenocytes from CD8-cre and CD8-cre *Nr4a1^fl/fl^ Nr4a3^−/−^* (cDKO) mice were stimulated with PMA and ionomycin (PMA/Io) for 2 h. Representative flow plots showing intra-cellular NUR77 expression following fixation and permeabilization within thymic and splenic T cell subsets. Plots are representative of at least 3 biological replicates. G. Quantification of NUR77 MFI in T cell subsets as depicted above in F (n = 3 biological replicates). H. Quantification of frequency of splenic CD8^+^CD44^hi^ T cells from CD8-cre, *Nr4a3^−/−^*, CD8-cre *Nr4a1^fl/fl^* and CD8-cre cDKO mice (n = 3 biological replicates). Graphs depict mean +/- SEM. Statistical significance was assessed by one-way (B, H) or two-way ANOVA with Dunnett’s test (D, E). *P < 0.05; ***P < 0.001; ****P < 0.0001. NS, not significant.

However, despite restoration of a replete Treg compartment of WT origin, DKO:WT chimeras nevertheless exhibit marked accumulation of CD44^hi^CD8^+^ T cells relative to WT:WT control chimeras (**Figures 5C, D**), and moreover, DKO T cells accumulate in this compartment (**Figure 5E)**. In addition, these CD44^hi^CD8^+^ DKO T cells upregulate PD-1 expression, suggesting an exhausted state (**Supplementary Figures 5C, D**). These observations reveal a T cell-intrinsic role for the NR4A family in CD8^+^ T cell homeostasis, and suggest self-reactive DKO CD8^+^ T cells may encounter chronic antigen stimulation in the periphery.

### Abnormal DKO CD8^+^ T cell homeostasis in mixed chimeras is due to a cell-intrinsic role for *Nr4a1* and *Nr4a3* during thymic development

To test whether abnormal DKO CD8^+^ T cell homeostasis reflects a requirement for the NR4A family during thymic development or exclusively in the periphery, we took advantage of a CD8-cre construct driven by the E8I enhancer that expresses specifically in mature CD8SP and peripheral CD8^+^ T cells in order to generate CD8-cre *Nr4a1^fl/fl^ Nr4a3^−/−^* mice (CD8-cre cDKO) (46) (**Figures 5F, G**). We can confirm that this cre is not active until after thymic DP stage and positive selection checkpoint are traversed, because NUR77 expression in the mature CD4 lineage of CD8-cre cDKO mice is intact (**Figures 5F, G**). Since accumulation of CD44^hi^CD8^+^ T cells was not observed in CD8-cre cDKO mice (**Figure 5H, Supplementary Figure 5E**), we conclude that this phenotype must be attributable to a role for the NR4A family earlier in development and likely reflects escape of self-reactive CD8^+^ T cells into the periphery due to impaired negative selection.

### Cell-intrinsic accumulation of CD4^+^ DKO T cells with anergic phenotype in mixed chimeras

gDKO mice exhibit an expanded CD4^+^ T cell effector-memory compartment that is not evident in DKO:WT chimeras (**Figures 6A-C**). However, CD4^+^ T cell homeostasis is not restored in these chimeras; rather DKO CD4^+^ T cells accumulate in the CD44^hi^ (memory) compartment and upregulate well-established markers of anergy (CD73 and FR4) in a cell-intrinsic manner (**Figures 6D-G**) (47, 48). Similarly, expansion of anergic CD4^+^ T cells was exaggerated in 1:1 DKO:WT chimeras relative to both control chimeras and SKO mice (**Supplementary Figures 6A-H**). Moreover, even phenotypically ‘naïve’ DKO CD44^lo^CD62L^hi^ CD4^+^ T cells in mixed chimeras acquired an anergic phenotype, suggestive of antigen encounter (**Figure 6D, E, Supplementary Figures 6A, C, F**) (26). Taken together, these data are consistent with escape of self-reactive DKO CD4^+^ T cells from negative selection in the thymus (**Figure 2**) and acquisition of an anergic phenotype in the periphery.

**Figure 6.**
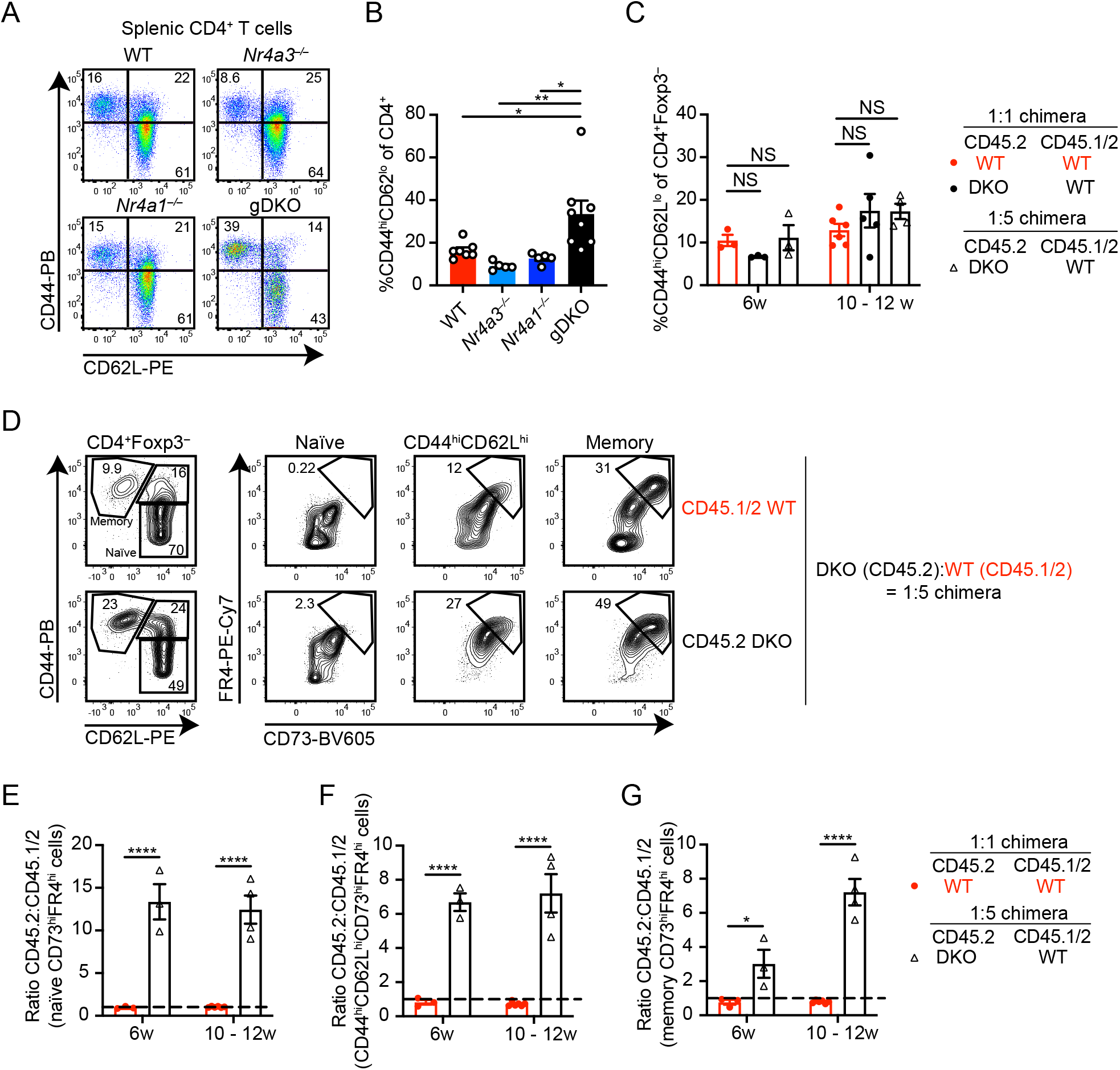
Accumulation of DKO CD4^+^ T cells with anergic phenotype in chimeras. A. Splenocytes from WT, *Nr4a3^−/−^*, *Nr4a1^−/−^* and gDKO mice were stained to detect CD4^+^ T cell subsets on the basis of CD44 and CD62L expression. Plots are representative of at least 5 mice. B. Quantification of splenic CD4^+^ CD44^hi^CD62L^lo^ T cells as gated in A (n ≧ 5 biological replicates, 3 to 4-week-old gDKO and 5 to 6-week-old mice with the other genotypes were analyzed). C. Quantification of splenic CD4^+^ CD44^hi^CD62L^lo^ T cells (after exclusion of FOXP3^+^ Treg) as gated in Supp. Fig. 6A from competitive chimeras at time points indicated following irradiation and BM transfer (n ≧ 3 biological replicates). D. Splenocytes from 12 weeks post-transplant DKO:WT = 1:5 chimera were stained with CD4, CD44, CD62L, CD73, FR4, CD45.1 and CD45.2, then permeabilized and stained with FOXP3. Shown are representative flow plots to detect CD73^hi^FR4^hi^ (anergic) T cells within CD44^lo^CD62L^hi^ (naïve), CD44^hi^CD62L^hi^ and CD44^hi^CD62L^lo^ (memory) compartments of CD4^+^FOXP3^−^ cells of each donor genotype. Plots are representative of 3 mice. E-G. Ratio of CD45.2 to CD45.1/2 within CD73^hi^FR4^hi^ gate among naïve (E), CD44^hi^CD62L^hi^ (F) or memory (G) CD4^+^ T cell compartments, as gated in D. Shown are WT:WT = 1:1 and DKO:WT = 1:5 chimeras at indicated time points following irradiation (n ≧ 3 biological replicates). Graphs depict mean +/- SEM. Statistical significance was assessed by one-way (B) ANOVA with Tukey’s test, two-way ANOVA with Dunnett’s test (C) or two-tailed unpaired Student’s t-test with the Holm–Šídák method (E-G). *P < 0.05; **P < 0.01; ****P < 0.0001. NS, not significant.

### Impaired proximal TCR signaling in anergic DKO T cells from mixed chimeras

Canonical functional features of anergic T cells include defective proximal TCR signal transduction and impaired IL-2 production (49, 50). Therefore, we next assessed TCR-induced Erk phosphorylation in T cells from DKO chimeras via a well-established flow-based assay. Since anergic surface markers were largely preserved after TCR stimulation and methanol permeabilization, we were able to gate cells according to CD73 and FR4 expression, and on this basis defined cells as non-anergic, intermediate anergic, or anergic (**Supplementary Figure 7A**). Consistent with this surface phenotype, we observed progressive hypo-responsiveness of WT memory CD4^+^ T cells across these populations (**Figures 7A, B**). Within each gate, DKO CD4^+^ T cells were even more refractory to TCR stimulation than WT cells from the same chimera. Strikingly, unlike cells of WT origin, naïve anergic DKO T cells were as refractory as memory anergic T cells. These data suggest that DKO CD4^+^ T cells not only upregulate markers of anergy (**Figure 6**), but acquired functional features of anergy and did so to an even greater extent than WT.

**Figure 7.**
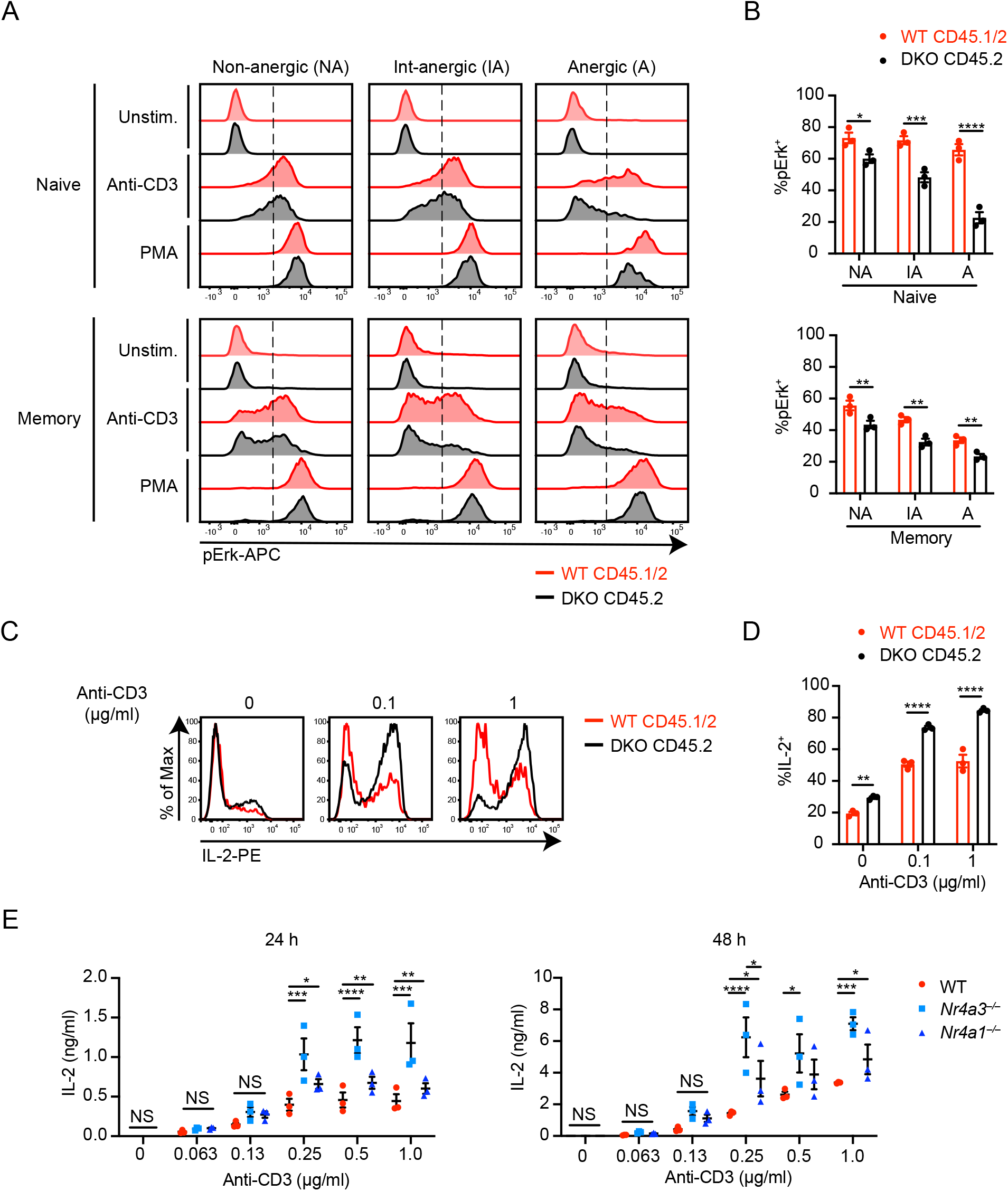
Cell-intrinsic defect in peripheral CD4^+^ T cell tolerance in DKO competitive chimeras. A. Splenocytes from DKO:WT = 1:5 chimera were stimulated with 10 μg/ml of anti-CD3 for 30 seconds followed by 50 μg/ml of secondary crosslinking antibody for 2 minutes, or alternatively with PMA for 2 minutes. Cells were fixed, permeabilized, and then stained with CD4, CD8, CD44, CD45.1, CD45.2, CD62L, CD73, FOXP3, FR4 and pErk. Representative histograms showing intra-cellular pErk expression in non-anergic (CD73^lo^FR4^lo^; NA), int-anergic (CD73^int^FR4^int^ ; IA) or anergic (CD73^hi^FR4^hi^; A) among naïve (CD44^lo^CD62L^hi^) or memory (CD44^hi^CD62L^lo^) CD4^+^ T cells gated as Supp Fig. 7A. Vertical dashed line shows the threshold of positive gate. Plots are representative of n = 6 mice. B. Quantification of %pErk^+^ as shown in A above (n = 3 biological replicates, representative of n = 2 independent experiments). C. Lymph node cells from 10 weeks post-transplant DKO:WT = 1:1 chimera were cultured in plates coated with indicated doses of anti-CD3 for 20 h. Then PMA, ionomycin and brefeldin were added and cells were cultured for additional 4 h. Cells were permeabilized and stained for CD4, CD8, CD45.1, CD45.2 and IL-2. Representative histograms showing intracellular IL-2 in CD4^+^ cells of each donor genotype. Plots are representative of 3 mice. D. Quantification of %IL-2^+^ as described for C above (n = 3 biological replicates). E. CD4^+^ cells were isolated by negative selection from lymph nodes and cultured in plates coated with indicated dose of anti-CD3 + 2 μg/ml anti-CD28 for 24 h (left) or 48 h (right). IL-2 concentration of supernatant was measured with ELISA (n = 3 biological replicates). Graphs depict mean +/- SEM. Statistical significance was assessed by two-tailed unpaired Student’s t-test with the Holm–Šídák method (B, D) or two-way ANOVA with Tukey’s test (E). *P < 0.05; **P < 0.01; ***P < 0.001; ****P < 0.0001. NS, not significant.

Since we observed the accumulation of DKO CD44^hi^CD8^+^ T cells with increased PD-1 expression in DKO:WT chimeras (**Figure 5, Supplementary Figures 5C, D**), we utilized the same approach to assess functional characteristics of DKO CD8^+^ T cells. We found that DKO CD44^hi^CD8^+^ T cells were functionally hyporesponsive relative to WT cells within the same chimera (**Supplementary Figures 7B-D**).

Of note, Erk phosphorylation downstream of PMA stimulation was intact in both genotypes across all gated populations, suggesting a proximal defect in TCR signaling among ‘tolerant’ T cells (**Figure 7A, Supplementary Figure 7C**). Importantly, TCR-induced Erk phosphorylation was robust in naïve / non-anergic DKO T cells (**Figures 7A, B, Supplementary Figures 7C, D**), implying that defective signal transduction was an acquired feature of tolerant T cells. Collectively, these data suggest that self-reactive DKO T cells escape negative selection and not only acquire phenotypic features of antigen experience, but also become hyporesponsive to TCR stimulation.

### Cell-intrinsic defect in peripheral CD4^+^ T cell tolerance in DKO competitive chimeras

Although we find that DKO CD4^+^ T cells acquire phenotypic and functional features of anergy, *Nr4a1* has recently been implicated in mediating CD4^+^ T cell tolerance (24). Indeed, a related study revealed that triple knockout (TKO) CD8^+^ chimeric antigen receptor (CAR) T cells lacking all *Nr4a* expression evaded exhaustion and downregulated anergy-related genes (17). One of many critical target genes identified in these studies was IL-2; impaired IL-2 production is among the most characteristic features of anergic T cells, while exogenous IL-2 can override anergy in some settings (50, 51). We therefore sought to assess IL-2 responses by DKO T cells. To do so, we cultured T cells from DKO:WT chimeras with anti-CD3 and then assessed the capacity for IL-2 production following maximal restimulation with PMA/ionomycin. We reasoned that this ought to unmask differences in TCR-induced chromatin remodeling around the IL-2 locus in DKO and WT T cells. We observed that, after TCR stimulation, DKO CD4^+^ T cells acquire a much higher capacity for IL-2 production relative to WT, and this is cell-intrinsic (**Figures 7C, D**). This result is not due to Treg deficiency in DKO compartment (**Supplementary Figure 7E**). *Nr4a1^−/−^* or *Nr4a3^−/−^* SKO CD4 T cells each exhibit a less robust but independent increase in capacity for IL-2 production relative to WT (**Supplementary Figure 7F**), as well as enhanced IL-2 secretion across a broad titration of TCR stimulation (as measured in culture supernatants via ELISA) (**Figure 7E**). These data suggest that the role of the NR4A family in restraining IL-2 production is not completely redundant and impacts naïve as well as anergic CD4^+^ T cells.

DKO CD8^+^ T cells also exhibited a much higher capacity for IL-2 production than WT cells from the same mixed chimera (**Supplementary Figures 7G, H**). Furthermore, CD8^+^ T cells from CD8-cre cDKO mice exhibit a nearly identical phenotype that is more robust than in SKO CD8^+^ T cells from *Nr4a3^−/−^* or CD8-cre *Nr4a1^fl/fl^* mice (**Supplementary Figure 7I**). These data suggest that the NR4A family negatively regulate the IL-2 locus in peripheral CD8^+^ T cells in a manner that is additive and cell-intrinsic, and independent of self-reactivity.

### Restoring Treg compartment in competitive chimeras alters autoantibody repertoire but does not suppress autoimmunity

gDKO mice exhibit spontaneous, early-onset development of autoantibodies (**Figures 8A, B**). Indirect immunofluorescence assay (IFA) for autoantibodies revealed both nuclear and cytosolic staining suggesting a widespread loss of B cell tolerance that occurs with complete penetrance before 4 weeks of age, recapitulating observations in Treg-deficient mice (8, 52–54). It is possible that this is attributable in part to loss of T follicular regulatory (Tfr) cells in gDKO as seen in other Treg-deficient mice (55, 56). Indeed, we identify a profound cell-intrinsic defect in Tfr (but not Tfh, as previously reported (57)) generation in DKO chimeras (**Supplementary Figures 8A, B**). Although older *Nr4a3*^−/−^ (but not *Nr4a1*^−/−^) mice exhibit very low titer autoantibodies with a similar pattern (**Supplementary Figures 8C, D**), B cell tolerance is largely preserved in SKO mice. To our surprise, despite reconstitution of the Treg (and Tfr) compartment in DKO:WT chimeras with cells of WT origin, we nevertheless observed the development of high titer autoantibodies even at early time points after reconstitution (**Figures 8C, D**). Cytosolic staining by autoantibodies was largely eliminated in these chimeras, but anti-nuclear autoantibodies persisted (**Figure 8E**). Consistent with this finding, anti-dsDNA antibodies were detected in 1:1 DKO:WT chimera but not in WT:WT chimera (**Figure 8F**). Importantly, we observe the development of ANA even in 1:5 DKO chimeras (**Supplementary Figures 8E, F**). This suggests that the development of autoimmunity in DKO chimeras was not attributable to a residual or partial Treg defect. The titer of ANA in DKO:WT chimeras increased with age (**Figure 8G, Supplementary Figure 8E**). This correlated with progressive accumulation of anergic CD4^+^ T cells, thymic atrophy, and development of polyclonal B cell activation and spontaneous GC expansion, suggestive of evolving immune dysregulation in these chimeras. By contrast, mice in which B cells conditionally lack both *Nr4a1* and *Nr4a3* (mb1-cre cDKO) did not develop ANA even after 40 weeks (**Supplementary Figures 8G, H**). These data imply that ANA appearing in DKO:WT chimeras are not attributable to a B cell-intrinsic role for the NR4A family. Rather, we propose that - while restoring Treg compartment suppresses lethal immune dysregulation in DKO chimeras - tolerance is not fully restored, and this may be due to a profound defect in both negative selection and peripheral T cell tolerance.

**Figure 8.**
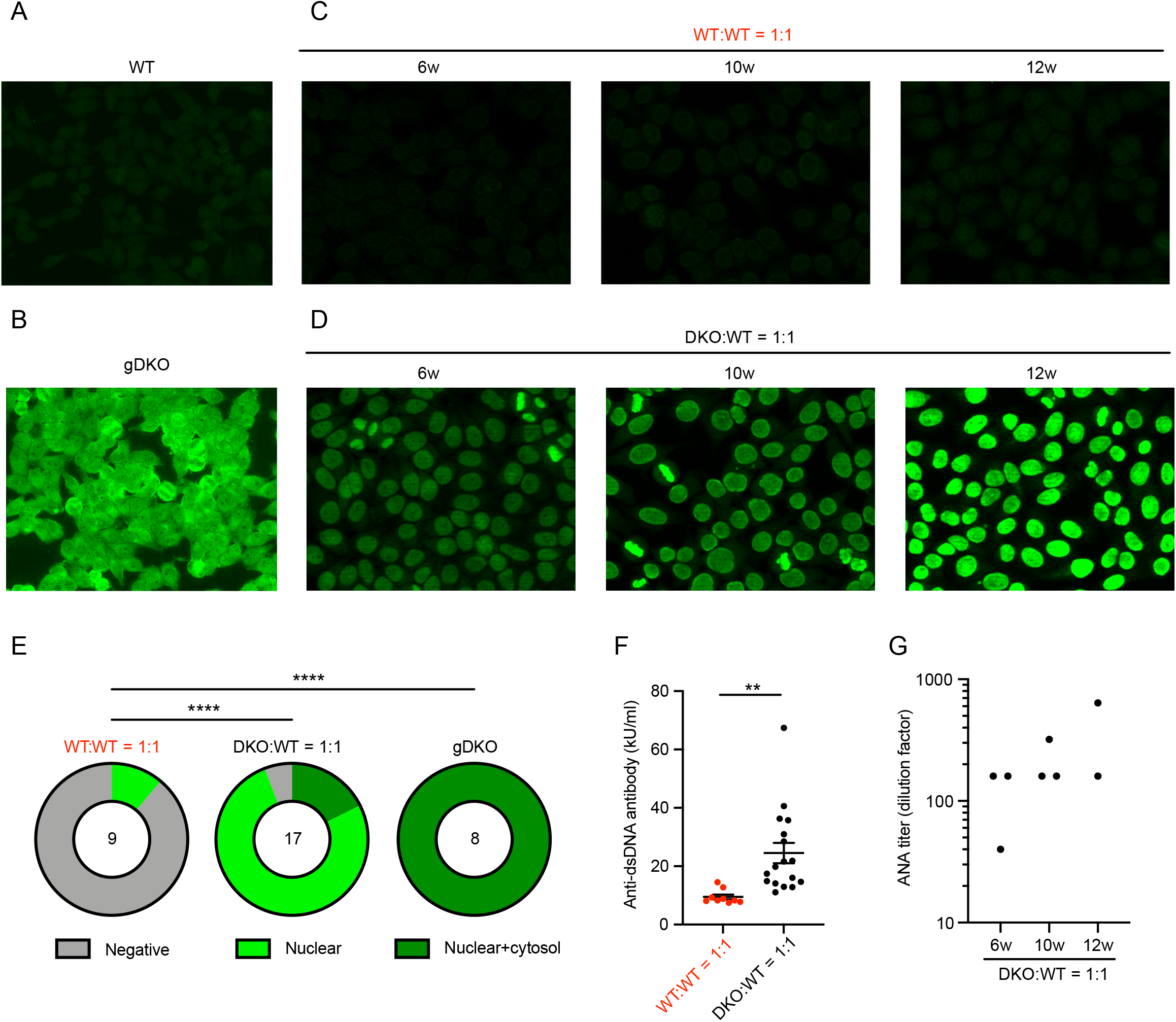
Restoring Treg compartment in competitive chimeras alters autoantibody repertoire but does not restore tolerance. A-D. Anti-nuclear antibody (ANA) immunofluorescence images. 1:40 diluted serum of indicated mice were applied to Hep-2 ANA substrate slides. Slides were stained with FITC-anti mouse IgG after washing. Images are representative of biological replicates as quantified below in E, G. E. Graphs depict frequency of negative, nuclear, or nuclear+cytoplasmic Hep-2 cell staining patterns in WT:WT = 1:1 chimera, DKO:WT = 1:1 chimera and gDKO. Statistical significance comparing ANA positivity was assessed by Fisher’s exact test. ****P < 0.0001. F. Quantification of anti-dsDNA antibody in serum from WT:WT = 1:1 chimera and DKO:WT = 1:1 chimera determined by ELISA for n = 9 (WT:WT = 1:1 chimera) and n = 17 (DKO:WT = 1:1 chimera) biological replicates. Statistical significance was assessed by two-tailed unpaired Student’s t-test. **P < 0.01. G. ANA titer determined with serial dilution of serum from chimeras at indicated time points post-transplant. Serum was serially diluted 2-fold from 1:40 to 1:1280. Hep-2 ANA substrate slides were stained with diluted serum. A blinded rheumatologist read the images and determined titer as the lowest detectable dilution of each sample.

## DISCUSSION

A vital and redundant role for NR4A factors in the Treg compartment has made it challenging to isolate and dissect other functions for this family in immune tolerance and homeostasis. Unfortunately, conditional genetic strategies alone cannot disentangle requirements for the NR4A family during thymic selection from their obligate function in Treg. Here we used competitive BM chimeras to reconstitute a functional Treg compartment and this enabled us to unmask additional essential roles for the NR4A family in the preservation of both central and peripheral T cell tolerance under homeostatic conditions.

We confirmed a cell-intrinsic requirement for *Nr4a1* and *Nr4a3* in the Treg compartment as previously reported with CD4-cre conditional DKO and TKO mice (14, 15, 58, 59). Concurrently, we also observed cell-intrinsic expansion of DKO CD25^+^ FOXP3^−^ cells both in gDKO mice and DKO chimeras. We speculate these cells represent either ‘ex-Treg’ that have lost FOXP3 expression, or cells that failed to differentiate into functional Treg, but favor the latter as this population is equally apparent in thymus and periphery (34) (**Figures 1K, O**). Sekiya and colleagues propose that this compartment contains highly self-reactive T cells that failed to assume Treg fate and yet escaped censorship by negative selection (15). This population of cells may contribute to the tolerance break we observed in DKO chimeras, but importantly, cells diverted from the Treg fate cannot account numerically for excess DKO SP thymocytes that escape negative selection in mixed chimeras, especially since equal or greater advantage for DKO cells relative to WT is observed in CD8SP thymocytes relative to CD4SP.

By contrast, we showed that myeloid cell expansion in gDKO mice is not cell-intrinsic because it is almost entirely suppressed in DKO chimeras, and is instead likely attributable to loss of Treg as observed in other Treg-deficient mouse models (6, 9, 42, 43). In support of this hypothesis, CD4-cre TKO mice develop a similar myeloproliferative disorder that is also rescued in competitive chimeras generated with mixtures of CD4-cre TKO and WT donor BM (14). It has been reported that expansion of CD11b^+^F4/80^+^ monocytes/macrophages in Treg-deficient Scurfy mice may be driven by high levels of M-CSF and GM-CSF (9, 42). We speculate that a similar phenomenon accounts for the myeloproliferative disorder seen in gDKO mice. This result emphasizes the need to critically re-assess the therapeutic potential of the NR4A family as drug targets in myeloproliferative disorders.

One of the earliest functions identified for the NR4A family is an essential role in Ag-induced cell death in T cell hybridomas (38, 39). The NR4A family has also been implicated in thymic negative selection, but these studies have largely relied on mis-expression of full length and truncated NR4A Tg constructs under the control of the proximal Lck promoter (which is active early during the DN stage of thymic development) (20, 28, 29). By contrast, studies of *Nr4a1^−/−^* mice have revealed subtle phenotypes, consistent with redundancy among family members (30, 31, 60). Here we were able to unmask the redundancy between *Nr4a1* and *Nr4a3* during thymic negative selection in a physiological setting for the first time. It is estimated that approximately 1.7x10^7^ cells are deleted per day via negative selection in a mouse (61). This is six times more than the number of positively selected thymocytes that emerge in the same time frame. Since both 1:1 and 1:5 DKO chimeras harbor 4- to 6-fold more CD4SP and CD8SP DKO cells relative to WT cells when normalized to the pre-selection DP compartment (**Figure 2E**), we propose that NR4A-dependent deletion may account for most or all negative selection.

It has been argued that negative selection by ubiquitously-expressed self-antigens occurs at the DP stage in the thymic cortex and may account for 75% of all deletion (61), while negative selection by tissue-restricted antigens (TRA) displayed on mTEC occurs at the early SP stage after CCR7 upregulation and migration to the thymic medulla (62). Prior studies of truncated dominant-negative NR4A Tgs and *Nr4a1^−/−^* mice in combination with H-Y and OTI/OTII RIP-mOVA models implicate the NR4A family in both processes (29, 31, 63–65). Though our data does not directly distinguish between the two, the striking amplitude of rescue seen in DKO chimeras suggests escape from negative selection by ubiquitous self-antigens and possibly TRA as well. The role of NR4A family remains to be tested in the context of DKO or TKO cells coupled with physiologically expressed TCR transgenes directed against either ubiquitous antigens or TRAs.

Caspase-3 is activated in thymocytes upon TCR stimulation and in the process of negative selection (66). Reduced aCasp3 expression of *in vitro* stimulated DKO thymocytes suggests caspase-dependent TCR-induced apoptosis is mediated, at least in part, by the NR4A family. BIM/*Bcl2l11*, a member of Bcl-2 family that can promote Caspase-3 activation is also essential for thymic negative selection and may represent a transcriptional target for *Nr4a1* (31, 67). Although it is possible that *Nr4a1* and *Nr4a3* might redundantly regulate *Bcl2l11* transcription, it has also been shown that NUR77 can promote apoptosis by directly binding BCL-2 in the cytosol, inducing a conformational change that exposes its BH3 pro-apoptotic domain in a manner that is independent of transcriptional activity of NUR77 (64, 68). It is important to further define how this pathway intersects with BIM-mediated negative selection, and whether there may be distinct upstream signals, developmental stages, or sites in the thymus where one or the other factor plays a dominant role, or whether instead classic genetic epistasis might define a linear pathway (60, 63).

It remains to be determined how additional instructional signals modulate NR4A function to promote Treg differentiation or, alternatively, drive deletion of self-reactive thymocytes. Factors such as IL-2 supply, co-stimulatory signals on distinct APC populations, or developmental context may be instructive (62). Another possible determinant is intensity/duration/biased signal transduction downstream of the TCR itself, which may in turn guide localization of NR4As to the cytosol or nucleus via post-translational modifications (69).

We observe the accumulation of DKO CD44^hi^CD8^+^ T cells in DKO competitive chimeras, and this is eliminated in CD8-cre cDKO mice in which cre-mediated deletion occurs only after thymic selection is complete (**Figure 5**). We also observed a marked accumulation of DKO cells in the CD73^hi^FR4^hi^ anergic CD4^+^ T cell compartment with associated suppression of proximal TCR signal transduction (**Figure 6, 7**). We propose that these phenotypes reflect escape of self-reactive T cells into the periphery due to a defect in central tolerance, and subsequent self-antigen encounter.

Recent work suggests that *Nr4a1* is required for induction and/or maintenance of CD4^+^ T cell anergy (24); over-expression of *Nr4a1* drives upregulation of a subset of anergy-related genes, whereas deletion of *Nr4a1* prevents generation of functionally tolerant T cells. Similarly, *Nr4a* TKO CAR T cells evade exhaustion and eliminate tumors (17). Although DKO T cells acquire features of tolerance in chimeras, we nevertheless observed the development of systemic immune dysregulation and ANA in DKO chimeras despite reconstitution of a functional Treg compartment, suggesting a residual defect in functional anergy. Indeed, although suppression of IL-2 production is among the most characteristic features of anergic T cells, we report enhanced capacity for IL-2 production in SKO T cells (consistent with prior studies of *Nr4a1^−/−^* T cells (24)), and much more so in DKO T cells. We suggest this reflects a role for the NR4A family in epigenetic remodeling of the *Il2* locus in response to TCR stimulation. Indeed, NR4A transcription factors modulate chromatin structure in the setting of chronic antigen engagement (17, 24), and interrogation of a recently published ATACseq data set reveals differentially accessible regions of open chromatin (OCR) near the *Il2* locus in *Nr4a3^−/−^* CD8^+^ T cells following 12 hr TCR stimulation (GSE143513) (70).

We initially hypothesized that autoantibody production in gDKO mice was largely mediated by loss of Treg (and Tfr) as in Scurfy and *Foxp3*-deficient mice (8, 52–54). Surprisingly, 1:1 and 1:5 DKO:WT competitive chimeras also developed high titer ANA at early time points and with high penetrance. These data argue strongly against a role for residual Treg dysfunction, as the Treg compartment in 1:5 chimeras is almost exclusively of WT origin. However, autoantibodies in chimeras are directed exclusively at nuclear autoantigens, and cytosolic Hep-2 staining pattern is largely suppressed, suggesting a role for Treg in suppressing a more widespread loss of B cell tolerance. By contrast, B cell-specific conditional DKO mice fail to develop ANA even after aging, implying a B cell-extrinsic role for the NR4A family in driving this phenotype. We propose that self-reactive DKO T cells that have escaped negative selection, Treg differentiation, and peripheral anergy accumulate in the periphery and drive ANA production in DKO chimeras. It remains to be defined whether defective central or peripheral tolerance (or both) are most relevant for the development of autoimmunity in DKO chimeras, and whether specific Th subsets (such as Tfh) play a role (59).

Nearly complete redundancy between *Nr4a1* and *Nr4a3* are evident in Treg and during negative selection; deletion of both family members is necessary to unmask these roles. By contrast, regulation of B cell responses (27) and CD8^+^ T cell exhaustion (17) by the NR4A family appear additive. Based on published work (24) and our observations of the IL-2 module in SKO and DKO T cells, we speculate that regulation of CD4^+^ T cell anergy is similarly additive, but this remains to be fully addressed. Although expression of *Nr4a2* is low in the T cell lineage under steady-state conditions, we also cannot exclude the possibility that *Nr4a2* compensates for and partially masks some immune phenotypes in DKO cells, especially in the context of inflammatory stimuli.

We propose that *Nr4a1* and *Nr4a3* regulate layered T cell tolerance mechanisms to preserve immune homeostasis *<u>under steady-state conditions</u>* (see model, **Supplementary Figure 9**). In addition, it is likely that NR4A factors also play important roles in counter-regulating acute as well as chronic inflammatory/immune stimuli and promoting a return to immune homeostasis. Indeed, negative feedback by NR4A restrain responses to LPS in myeloid cells (71) and to antigen stimulation in B cells (27). In fact, the NR4A family suppresses inflammation in the context of several immune-mediated disease models (72–74). These findings suggest that NR4A family members represent important therapeutic targets. Although it remains unclear whether endogenous ligands regulate NR4A function *in vivo*, small molecule NUR77 agonist (75) and antagonist (76) compounds have been reported. Agonists might be useful to suppress autoimmunity and maintain transplant tolerance by promoting thymic negative selection, differentiation of Treg, and induction and maintenance of peripheral T and B cell tolerance. Antagonizing NUR77 and perhaps other NR4A family members could have applications for cancer immunotherapy (17), and, as we previously proposed, as a universal adjuvant for T-independent vaccines (27). Since redundancy among NR4A family members is important in both negative selection and in Treg, selectively targeting individual NR4A family members may allow modulation of antigen-specific T and B cell responses without disrupting global immune homeostasis. Conversely, our studies unmask Treg-independent and redundant roles for *Nr4a1* and *Nr4a3* in maintaining T cell tolerance under homeostatic conditions, with important implications for drug design.

## MATERIALS AND METHODS

*<u>Mice.</u> Nr4a1^−/−^*, *Nr4a1^fl/fl^*, and *Nr4a3^−/−^* mice were previously described (14, 27, 30). *Nr4a1^fl/fl^* were previously obtained from Catherine Hedrick (La Jolla Institute for Immunology) with permission from Pierre Chambon (University of Strasbourg) (14). *Nr4a1^−/−^* mice were obtained from The Jackson Laboratory and this line is used throughout the manuscript exclusively as single germline knockout comparator (30), *Nr4a3^−/−^* mice were generated in our laboratory as previously described (27). CD8-cre and mb1-cre were obtained from The Jackson Laboratory (46, 77). C57BL/6 mice were from The Jackson Laboratory and CD45.1^+^ BoyJ mice were from Charles River Laboratories. To generate germline DKO *Nr4a1^−/−^ Nr4a3^−/−^* mice, we bred *Nr4a3^−/−^* and *Nr4a1^fl/fl^* mice with germline recombination of the loxp-flanked locus, and confirmed loss of exon 2 both by genomic DNA PCR and transcript qPCR. All strains were fully backcrossed to C57BL/6 genetic background for at least 6 generations. Mice of both sexes were used for experiments between the ages of 3 and 10 weeks except for BM chimeras as described below. All mice were housed in a specific pathogen-free facility at UCSF according to the University and National Institutes of Health guidelines.

### Antibodies and Reagents

#### Abs for surface markers

Abs to B220, CD3, CD4, CD8, CD11b, CD19, CD21, CD23, CD25, CD44, CD45.1, CD45.2, CD62L, CD69, CD73, CD86, CD93 (AA4.1), CD138, Fas, FR4, γδTCR, GL7, Gr1, IgD, MHC-II, NK1.1, PD-1, and pNK conjugated to fluorophores were used (BioLegend, eBiosciences, BD, or Tonbo).

#### Abs for intra-cellular staining

FOXP3 Ab conjugated to APC or FITC (clone FJK-16s) manufactured by Invitrogen, purchased from Thermo Fisher Scientific. Anti-active Caspase-3 (aCasp3) Ab conjugated to APC (clone C92-605) purchased from BD Pharmingen. Anti-NUR77 conjugated to PE (clone 12.14) was manufactured by Invitrogen and purchased from eBioscience. Anti-IL-2 Ab conjugated to PE (clone JES6-5H4) manufactured by Invitrogen, purchased from Thermo Fisher Scientific. Anti-pERK (Phospho-p44/42 MAPK (T202/Y204) (clone 197G2) Rabbit Ab was purchased from Cell Signaling Technologies. APC Goat Anti-Rabbit IgG (H+L) was purchased from Jackson ImmunoResearch.

#### Stimulatory Abs

Anti-CD3 (clone 2c11) and anti-CD28 (clone 37.51) were from BioLegend. Goat anti-Armenian Hamster antibody was from Jackson ImmunoResearch.

#### ELISA reagents

96-well, high-binding, flat-bottom, half-area, clear polystyrene Costar Assay Plate (Corning). Mouse anti-dsDNA IgG-specific ELISA Kit was from Alpha diagnostic. Mouse IL-2 DuoSet ELISA DuoSet and Ancillary Reagent Kit 2 were from R&D Systems.

#### Anti-nuclear antibody (ANA)

NOVA Lite^TM^ HEp-2 ANA Substrate Slide and mounting medium (INOVA Diagnostics, Inc, #708100); FITC donkey anti-mouse IgG (Jackson ImmunoResearch, #715-095-150).

#### Media

Complete culture media was prepared with RPMI-1640 + L-glutamine (Corning-Gibco), Penicillin Streptomycin L-glutamine (Life Technologies), HEPES buffer [10mM] (Life Technologies), B-Mercaptoethanol [55mM] (Gibco), Sodium Pyruvate [1mM] (Life Technologies), Non-essential Amino acids (Life Technologies), 10% heat inactivated FBS (Omega Scientific).

#### Flow Cytometry

After staining, cells were analyzed on a Fortessa (Becton Dickson). Data analysis was performed using FlowJo (v9.9.6 or v10.7.1) software (Becton Dickson).

#### Intracellular staining to detect active Caspase 3

Following *in vitro* stimulation, cells were permeabilized and stained with APC-aCasp3, according to the manufacturer’s protocol (BD Cytofix/Cytoperm kit).

#### FOXP3 staining

FOXP3 staining was performed utilizing a FOXP3/transcription factor buffer set (eBioscience) in conjunction with APC or FITC anti-FOXP3, as per manufacturer’s instructions.

#### Intracellular staining to detect IL-2

Splenocytes were stimulated with plate-bound anti-CD3 Ab for 20 hours followed by a 4-hour treatment with 20 ng/ml of phorbol myristate acetate (PMA) (Sigma) and 1 μM of ionomycin (Calbiochem) and protein transport inhibitor cocktail (eBioscience) per manufacturer’s protocol. Following *in vitro* stimulation, cells were permeabilized and stained, according to the manufacturer’s protocol (BD Cytofix/Cytoperm kit).

#### Intracellular staining to detect NUR77

Following 2-hour *in vitro* stimulation with 20 ng/ml of PMA and 1 μM of ionomycin, cells were fixed in a final concentration of 4% paraformaldehyde for 10 min, permeabilized at −20 °C with 100% methanol for 30 min and, following washes and rehydration, stained with primary antibody for 60 min at 20 °C (room temperature).

#### Live/dead staining

LIVE/DEAD Fixable Near-IR Dead Cell Stain kit (Invitrogen). Reagent was reconstituted in DMSO as per manufacturer’s instructions, diluted 1:1000 in PBS, and cells were stained at a concentration of 1 × 10^6^ cells /100 μl on ice for 15 minutes.

#### Vital dye loading

Cells were loaded with CellTrace Violet (CTV; Invitrogen) per the manufacturer’s instructions except at 5 × 10^6^ cells/ml rather than 1 × 10^6^ cells/ml.

#### In vitro T cell culture and stimulation

Flat bottom 96 well plates were coated with varying doses of anti-CD3 with or without 2 mg/ml anti-CD28 at 4°C overnight. Splenocytes, lymphocytes or thymocytes were harvested into single cell suspension. Splenocytes were subjected to red cell lysis using ACK buffer. Cells were plated at a concentration of 5 × 10^5^ cells/100 μl complete RPMI media in antibody coated flat bottom 96 well plates for 1-4 days. In some cases, cells were loaded with CTV as described above prior to plating.

#### Bone marrow chimeras

Host mice were irradiated with two doses of 530 rads, 4 hours apart, and injected on the same day IV with a total of 2 × 10^6^ donor BM cells at varying ratios (1:1 or 1:5 or without mixture, as noted). Chimeras were sacrificed 6-14 weeks after irradiation for downstream analyses. *CD4^+^ T cell purification.* CD4^+^ T cell purification was performed utilizing magnetic-activated cell sorting (MACS) separation, per the manufacturer’s instructions. In brief, pooled spleen and/or lymph nodes were prepared utilizing the CD4^+^ T Cell Isolation Kit (Miltenyi Biotec) and purified by negative selection through an LS column (Miltenyi Biotec). Purified CD4^+^ T cells were then subjected to *in vitro* culture.

#### Phospho-flow

Splenocytes were rested at 37°C in serum-free RPMI for 30 minutes. Cells were then stimulated with 10 μg/ml of anti-CD3 (clone 2c11) for 30 seconds followed by 50 μg/ml of anti-Armenian hamster crosslinking antibody for 2 minutes, or PMA for 2 minutes. Stimulated cells were fixed with 2% paraformaldehyde and permeabilized with methanol at -20°C overnight. Cells were then stained with surface markers and pErk at 20°C.

#### ELISA for IL-2 detection

Purified lymph node CD4^+^ T cells were cultured on anti-CD3/28 coated plate at 1 × 10^5^ cells per well. Plates were spun and supernatants were harvested after 24h or 48 h. IL-2 concentrations in supernatants were measured using a commercial ELISA kit, per the manufacturer’s instructions (R&D Biosystems). In brief, 96-well plates were coated with 1 ug/ml of capture anti-IL-2 antibody. Supernatants were diluted serially, and the IL-2 was detected with detection anti-IL-2 antibody. ELISA plates were developed with mixture of tetramethylbenzidine and peroxidase, then stopped with 2 N sulfuric acid. Absorbance was measured at 450 nm using a spectrophotometer (SpectraMax M5; Molecular Devices).

#### ELISA for serum anti-dsDNA

Serum was harvested from blood collected by lateral tail vein sampling or cardiac puncture postmortem. Serum anti-dsDNA titer was measured with a commercial ELISA kit, per the manufacturer’s instructions (Alpha diagnostic). In brief, Sera were diluted serially and added to plates coated with dsDNA. Anti-dsDNA titer was detected with anti-IgG-HRP. ELISA plates were developed, and absorbance was measured as described above.

#### Anti-nuclear antibody (ANA)

Serum ANA was detected with NOVA Lite^TM^ HEp-2 ANA Substrate Slide as per manufacturer’s instructions except for using FITC-conjugated donkey anti-mouse IgG secondary antibody. Images were captured with a Zeiss Axio Imager M2 widefield fluorescence microscope. Images were processed with Zen Pro (Zeiss). To measure titer, serum was serially diluted 2-fold from 1:40 to 1:1280. HEp-2 ANA slides were stained with diluted serum. Images were read by a rheumatologist in a blinded manner and titer was determined as the detectable lowest dilution of each sample.

#### Statistical analysis

Statistical analysis and graphs were generated using Prism v9 (GraphPad Software, Inc). Graphs show mean ± SEM unless otherwise stated. Student’s unpaired or paired t-test was used to calculate the P values for all comparisons of two groups, and correction for multiple comparisons across time points or doses was then performed using the Holm–Šídák method. One-way or two-way analysis of variance (ANOVA) with follow-up Tukey’s test or Dunnett’s test were performed when more than two groups were compared with one another. Fisher’s exact test was used to compare the difference in proportions of two groups. *p<0.05, **p<0.01, ***p<0.001, ****p<0.0001.

### Author contributions

R.H., H.V.N., J.L.M. and J.Z. conceived of and designed the experiments. R.H., H.V.N. and J.L.M performed the experiments. R.H., H.V.N., J.L.M. and J.Z. analyzed the data. R.H. and J.Z. wrote the manuscript. R.H., H.V.N., J.L.M. and J.Z edited the manuscript.

## Acknowledgements

We thank Al Roque for help with mouse husbandry. We thank Arthur Weiss, Jeremy Brooks, Tran Nguyen, and Wan-Lin Lo for critical scientific feedback.

---

**Supplementary Figure 1. Supporting data for Figure 1.**
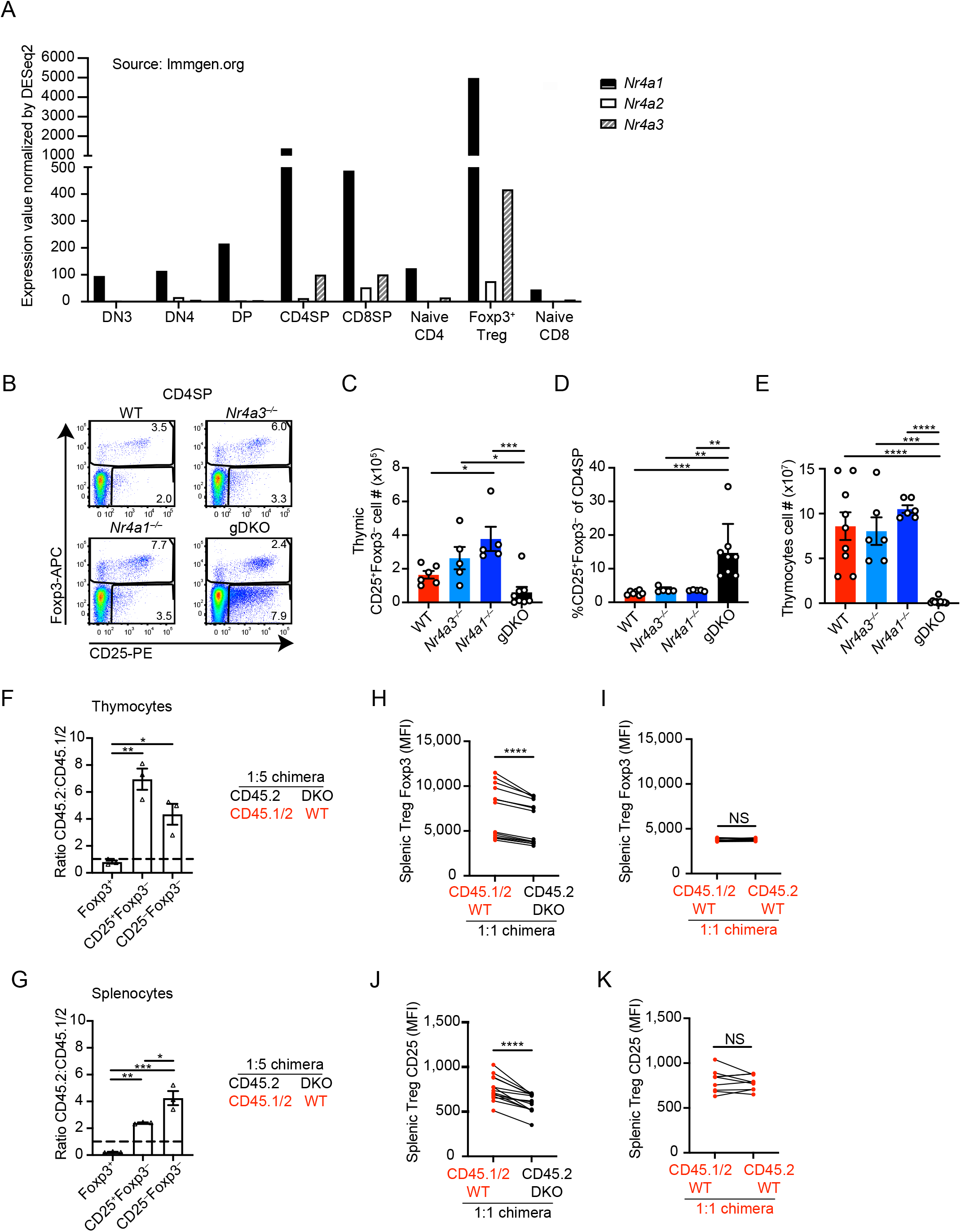
A. Expression of *Nr4a* family mRNA in T cell subsets quantified via RNA–seq. Data are exported from Immgen database (http://rstats.immgen.org/Skyline/skyline.html). B. Representative flow plots showing thymic Treg gate in WT, *Nr4a3^−/−^*, *Nr4a1^−/−^* and *Nr4a1^−/−^ Nr4a3^−/−^* (gDKO) mice, as determined by FOXP3 and CD25 expression. Plots are representative of at least 5 mice/genotype. C, D. Quantification of cell number (C) and frequency (D) of thymic CD4SP CD25^+^FOXP3^−^ cells as gated in (B). E. Quantification of viable thymocyte cell number from WT, *Nr4a3^−/−^*, *Nr4a1^−/−^* and gDKO mice. (Data in C-E include n ≧ 5 biological replicates/genotype, 3 to 4-week-old gDKO and 5 to 6-week-old mice with the other genotypes were analyzed). F, G. Ratio of CD45.2 to CD45.1/2 for Treg, CD25^+^FOXP3^−^ cells and CD25^−^FOXP3^−^ cells among thymic CD4SP (F) or splenic CD4^+^ (G) in DKO:WT = 1:5 chimera (n = 3 biological replicates). Ratios were normalized to DP thymocytes. H-K. Quantification of FOXP3 (H, I) or CD25 (J, K) expression (MFI) on splenic Treg cells of each donor genotype from DKO:WT = 1:1 (H, J) and WT:WT = 1:1 chimeras (I, K) determined via flow staining. Lines connect donor genotypes within an individual chimera (n = at least 9 biological replicates). Graphs depict mean +/- SEM. Statistical significance was assessed by one-way ANOVA with Tukey’s test (C-E, F, G) or two-tailed paired Student’s t-test (H-K). *P < 0.05; **P < 0.01; ***P < 0.001; ****P < 0.0001. NS, not significant.

**Supplementary Figure 2. Supporting data for Figure 2.**
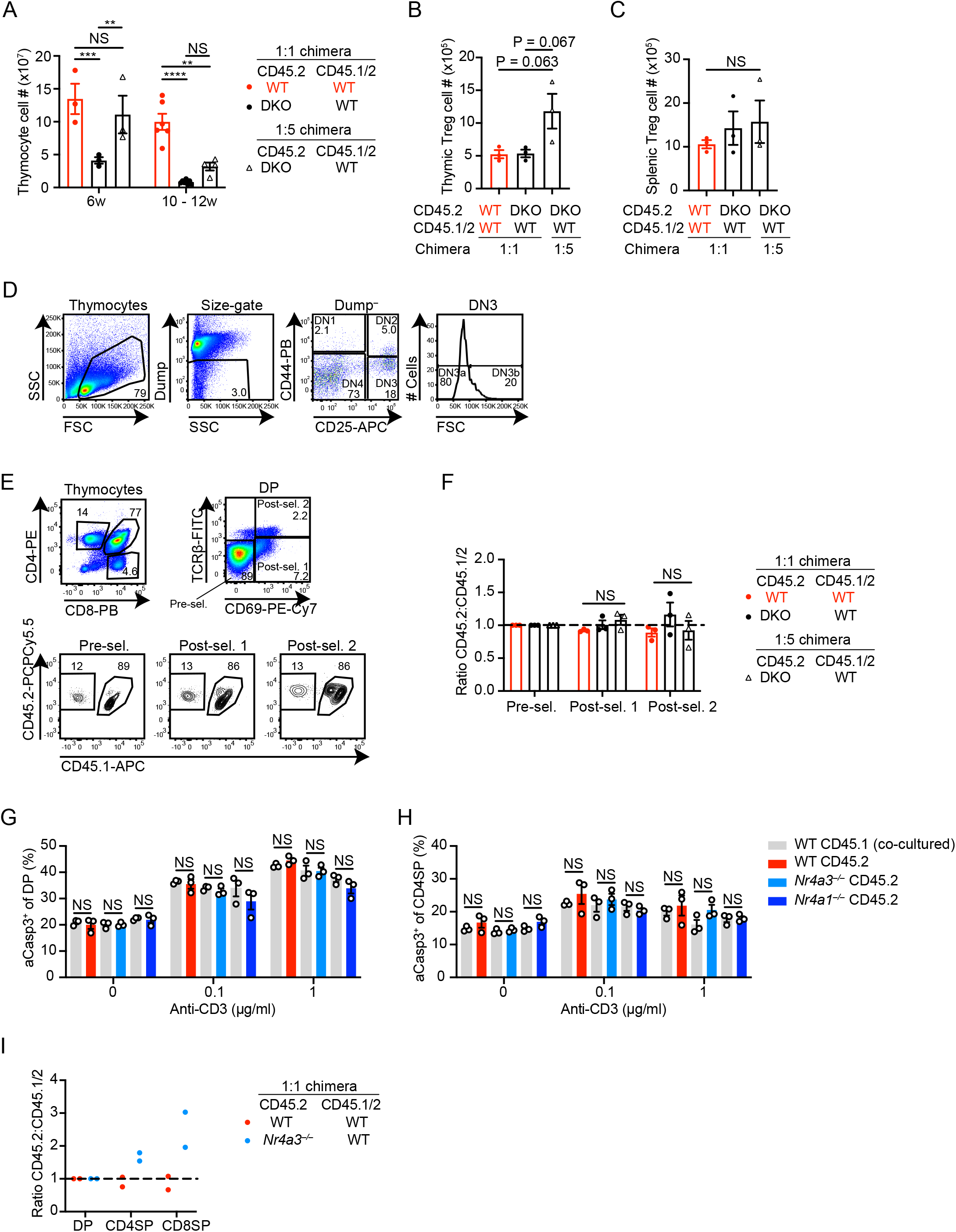
A. Quantification of thymocyte cell number in WT:WT = 1:1, DKO:WT = 1:1 and 1:5 chimera at indicated time points post-transplant (n ≧ 3 biological replicates). B, C. Quantification of thymic (B) and splenic (C) Treg cell number in WT:WT = 1:1, DKO:WT = 1:1 and 1:5 chimera at 6 weeks post-transplant (n = 3 biological replicates). D. Representative flow plots show gating strategy to identify pre- and post- b selection subsets DN3a/b in 1:1 DKO:WT chimera. Dump staining includes CD4, CD8, CD3, CD19, γδTCR, NK1.1, pNK, CD11b, Gr1, and CD11c. Plots are representative of 4 chimeras. E. Representative flow plots show gating strategy to detect pre- and post-positive selection DP subsets in DKO:WT = 1:5 chimera. Plots are representative of 3 chimeras. F. Ratio of CD45.2 to CD45.1/2 of pre-selection, post-selection 1 and post-selection 2 DP thymocytes (as gated in E) in chimeras, normalized to pre-selection DP (n = 3 biological replicates). G, H. Thymocytes from CD45.2 WT, *Nr4a3^−/−^* and *Nr4a1^−/−^* were mixed with CD45.1 WT thymocytes in 1:1 ratio. Cells were co-cultured, stimulated and stained as described for Figure 2F, I. Quantification of frequency of aCasp3^+^ cells among DP (G) or CD4SP (H) with indicated dose of anti-CD3 (n = 3 biological replicates). I. Ratio of CD45.2 to CD45.1/2 thymic subsets in WT:WT = 1:1 chimera and *Nr4a3^−/−^*:WT = 1:1 chimera, normalized to DP thymocytes (n = 2 biological replicates). Graphs depict mean +/- SEM. Statistical significance was assessed by two-way (A, F) or one-way (B, C) ANOVA with Tukey’s test or two-tailed unpaired Student’s t-test with the Holm–Šídák method (G, H). *P < 0.05; **P < 0.01; ***P < 0.001; ****P < 0.0001. NS, not significant.

**Supplementary Figure 3.**
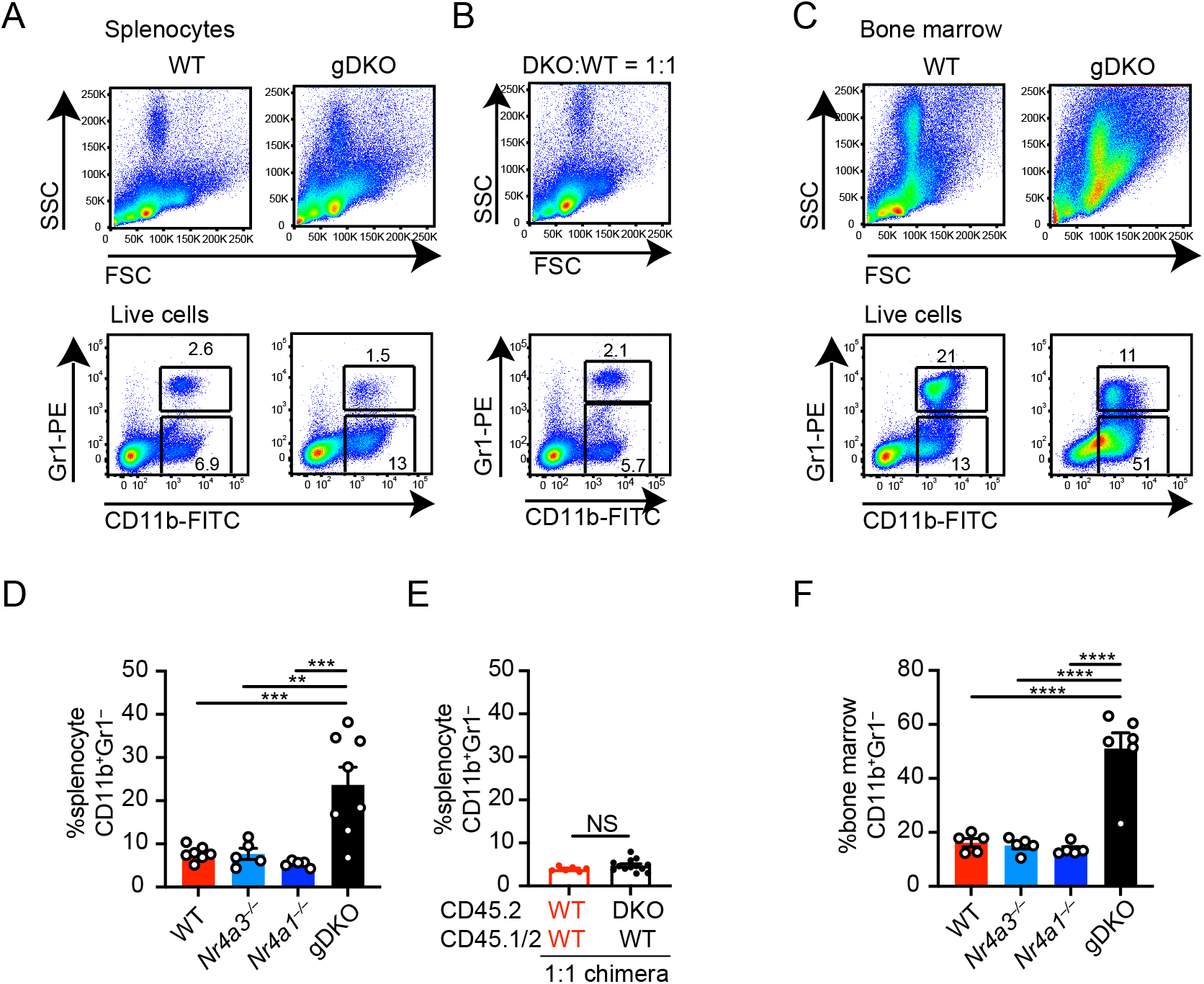
Expansion of myeloid cells in spleen and bone marrow of gDKO mice. A-C. Splenocytes (A) and BM cells (C) from WT and gDKO mice, as well as splenocytes from DKO:WT = 1:1 chimera (B) were stained as described for Figure 3A-D. Shown are representative plots of at least 5 mice. D, F. Quantification of CD11b^+^Gr1^−^ cell frequency in spleen (D) and bone marrow (F) from WT, *Nr4a3^−/−^*, *Nr4a1^−/−^* and gDKO mice (n ≧ 5 biological replicates, 3 to 4-week-old gDKO and 5 to 6-week-old mice with the other genotypes were analyzed). E. Quantification of CD11b^+^Gr1^−^ cell frequency in spleen from WT:WT = 1:1 and DKO:WT = 1:1 chimera (n ≧ 3 biological replicates). Graphs depict mean +/- SEM. Statistical significance was assessed by one-way ANOVA with Tukey’s test (D, F) or two-tailed unpaired Student’s t-test (E). **P < 0.01; ***P < 0.001; ****P < 0.0001. NS, not significant.

**Supplementary Figure 4. Supporting data for Figure 4.**
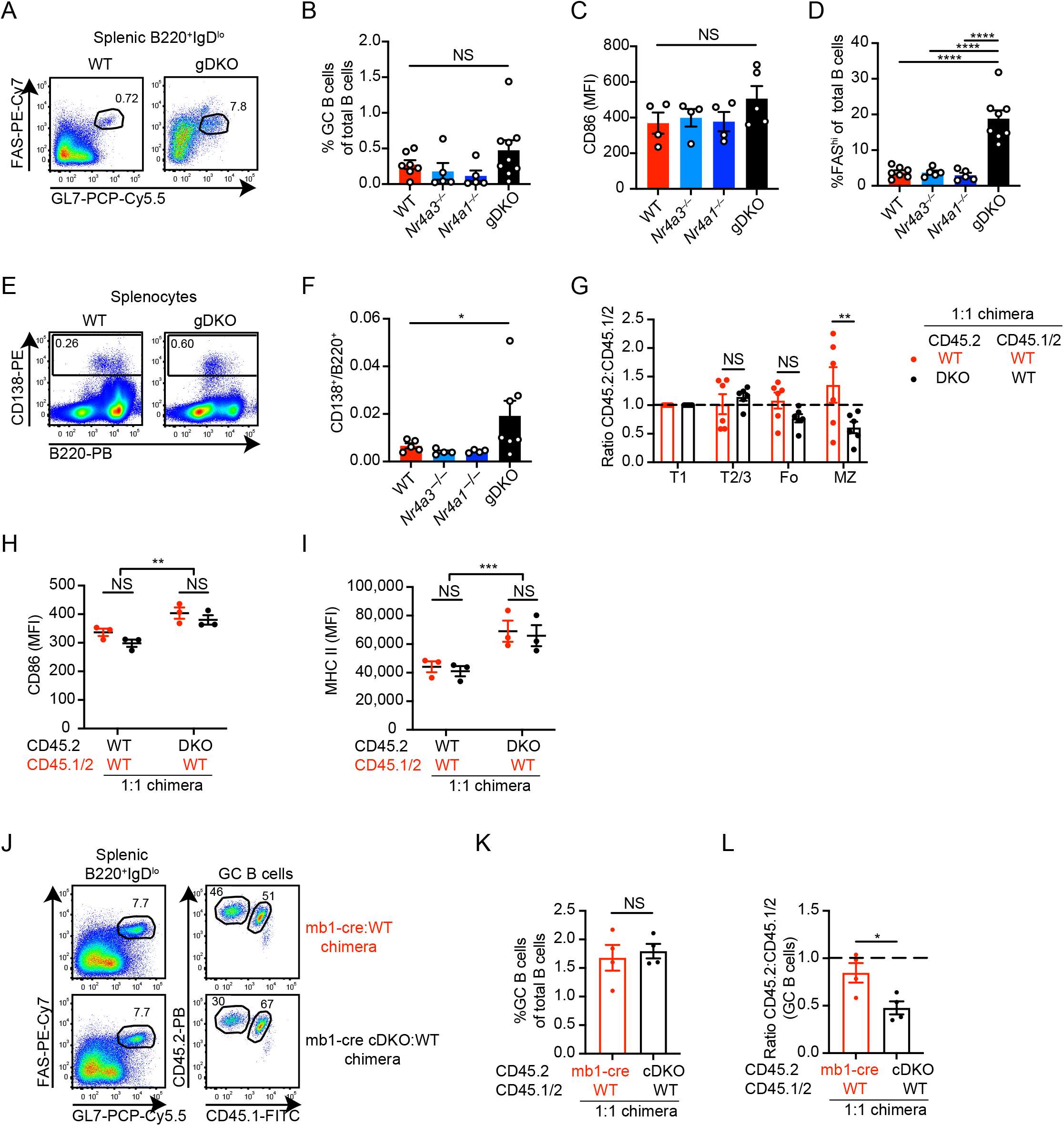
A. Representative flow plots show FAS^hi^GL7^+^ GC B cells from WT and gDKO mice as described for Figure 4D. Plots are representative of 6 mice. B. Quantification of frequency of GC B cells, as gated in (A) above, among total B cells from WT, *Nr4a3^−/−^*, *Nr4a1^−/−^* and gDKO mice (n ≧ 5 biological replicates). C. Quantification of CD86 MFI on splenic B cells from WT, *Nr4a3^−/−^*, *Nr4a1^−/−^* and gDKO mice (n ≧ 4 biological replicates). D. Quantification of FAS^hi^ cells among B220^+^ cells in spleen from WT, *Nr4a3^−/−^*, *Nr4a1^−/−^* and gDKO mice (n ≧ 5 biological replicates). E. Representative flow plots show CD138^+^ cells in splenocytes from WT and gDKO mice. Plots are representative of 4 mice. F. Quantification of ratio of CD138^+^ to B220^+^ cells, as gated in (E) above, from WT, *Nr4a3^−/−^*, *Nr4a1^−/−^* and gDKO mice (n ≧ 4 biological replicates). 3 to 4-week-old gDKO and 5 to 6-week-old mice with the other genotypes were analyzed (B, C, D, F). G. The ratio of CD45.2 to CD45.1/2 for transitional1 (T1), T2/3, follicular (Fo) and marginal zone (MZ) B cell subsets (gated on the basis of B220, CD21, CD23, and CD93) from WT:WT and DKO:WT = 1:1 chimera, normalized to T1 subset (n = 6 biological replicates). H, I. Quantification of CD86 (H) and MHC II (I) MFI on splenic B cells of each donor genotype in in WT:WT and DKO:WT = 1:1 chimera (n = 3 biological replicates). J. Representative flow plots showing FAS^hi^GL7^+^ GC B cells in spleen from mb1-cre:WT = 1:1 chimera and mb1-cre *Nr4a1^fl/fl^ Nr4a3^−/−^* (cDKO):WT = 1:1 chimera on the left. Right-hand plots depict CD45.2 cDKO donor and CD45.1/2 WT donor populations. Plots are representative of n = 4 biological replicates. K. Quantification of frequency of GC B cells among total B cells as described for J above (n = 4 biological replicates pooled from n = 2 independent experiments). L. Ratio of CD45.2 to CD45.1/2 GC B cells in spleen from chimeras described in (J) above, normalized to B220^+^IgD^hi^ cells (n = 4 biological replicates). Graphs depict mean +/- SEM. Statistical significance was assessed by Brown-Forsythe ANOVA (B, F), one-way ANOVA with Tukey’s test (C, D) or two-tailed unpaired Student’s t-test with (G, H, I) or without (K, L) the Holm–Šídák method. *P < 0.05; **P < 0.01; ***P < 0.001; ****P < 0.0001. NS, not significant

**Supplementary Figure 5. Supporting data for Figure 5.**
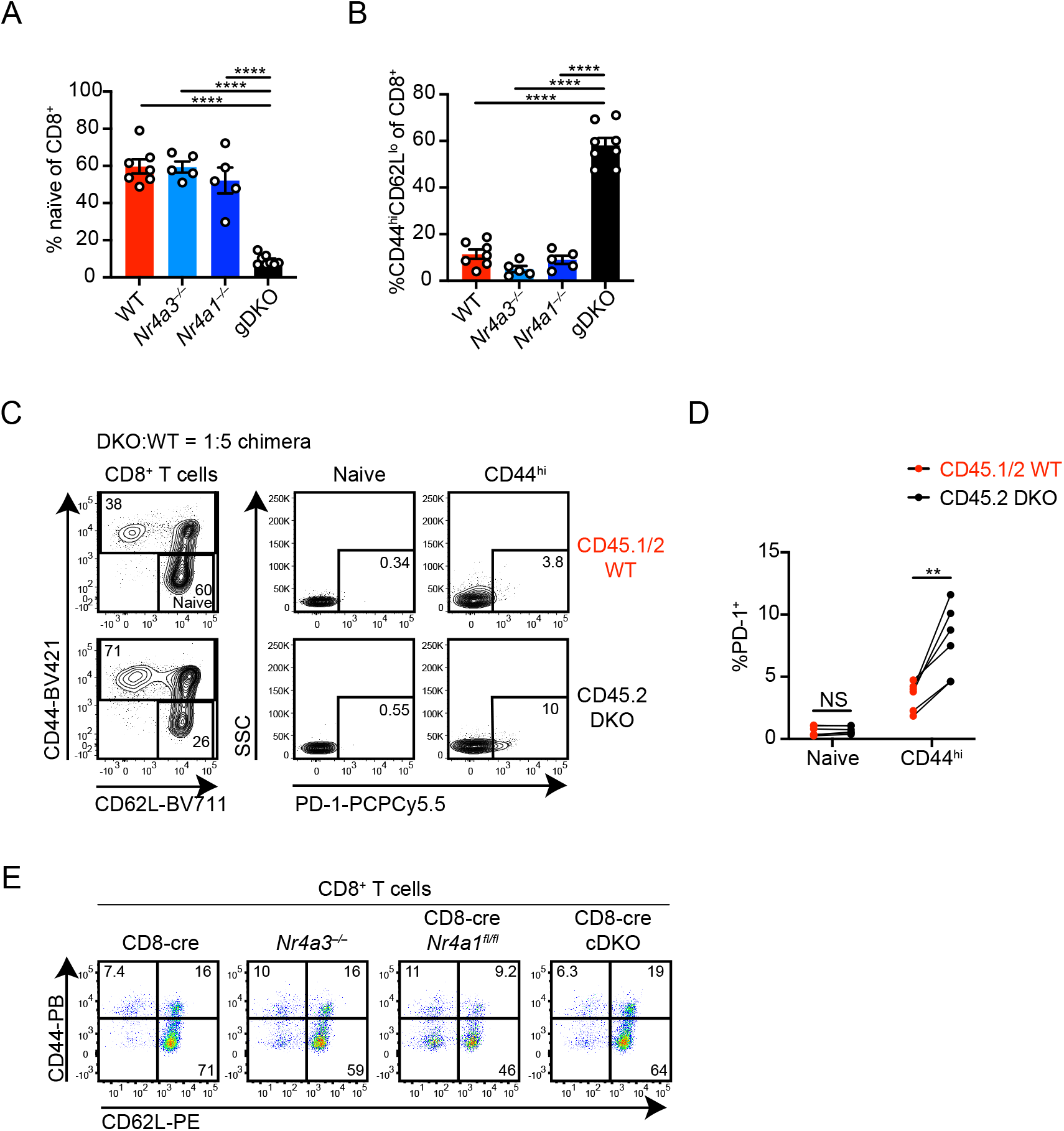
A, B. Quantification of frequency of CD8^+^ naïve CD44^lo^CD62L^hi^ (A) and CD44^hi^CD62L^lo^ (B) cells from WT, *Nr4a3^−/−^*, *Nr4a1^−/−^* and gDKO mice as gated in Fig. 5A (n ≧ 5 biological replicates, 3 to 4-week-old gDKO and 5 to 6-week-old mice with the other genotypes were analyzed). C. Splenocytes from DKO:WT = 1:5 chimera was stained to detect PD-1 expression on naïve CD44^lo^CD62L^hi^ and on CD44^hi^ CD8^+^ cells among each donor genotype. Plots are representative of 6 mice. D. Quantification of %PD-1^+^ of naïve and CD44^hi^ CD8^+^ cells as gated in C above. Lines connect donor genotypes within an individual chimera (n = 6 biological replicates). E. Splenocytes from CD8-cre, *Nr4a3^−/−^*, CD8-cre *Nr4a1^fl/fl^* and CD8-cre *Nr4a1^fl/fl^ Nr4a3^−/−^* (cDKO) mice were stained as described for Fig. 5A to detect CD8^+^ T cell subsets. Plots are representative of 3 mice and correspond to quantification in Fig. 5H. Graphs depict mean +/- SEM. Statistical significance was assessed by one-way ANOVA with Tukey’s test (A, B) or two-tailed unpaired Student’s t-test with the Holm–Šídák method (D). **P < 0.01; ****P < 0.0001. NS, not significant

**Supplementary Figure 6. Supporting data for Figure 6.**
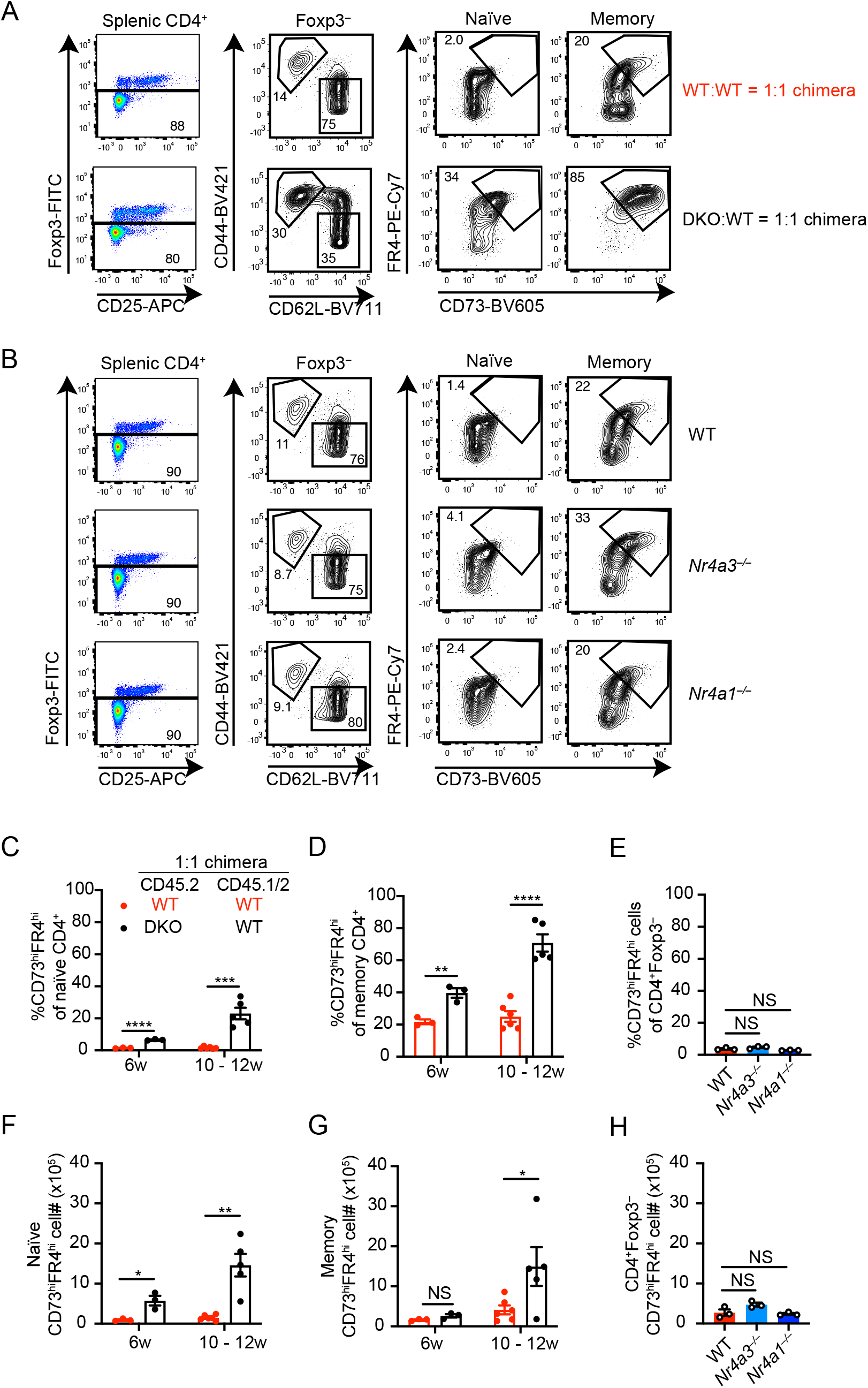
A. Splenocytes from 10 weeks post-transplant WT:WT = 1:1 and DKO:WT = 1:1 chimeras were stained as described for Fig. 6D to identify CD73^hi^FR4^hi^ (anergic) T cells. Plots are representative of 3 mice. B. Splenocytes from WT, *Nr4a3^−/−^* and *Nr4a1^−/−^* mice were stained as described for Fig. 6D. Plots are representative of 3 mice. C, D, F, G. Quantification of frequency (C, D) and cell number (F, G) of CD73^hi^FR4^hi^ T cells among naïve (C, F) or memory (D, G) CD4^+^ cells in WT:WT = 1:1 chimera and DKO:WT = 1:1 chimera at indicated time points post-transplant (n ≧ 3 biological replicates). E, H. Quantification of frequency (E) and cell number (H) of CD73^hi^FR4^hi^ T cells among total CD4^+^FOXP3^−^ T cells from WT, *Nr4a3^−/−^* and *Nr4a1^−/−^* mice (n = 3 biological replicates). Graphs depict mean +/- SEM. Statistical significance was assessed by two-tailed unpaired Student’s t-test with the Holm–Šídák method (C, D, F, G) or one-way ANOVA with Dunnett’s test (E, H). *P < 0.05; **P < 0.01; ***P < 0.001; ****P < 0.0001. NS, not significant

**Supplementary Figure 7. Supporting data for Figure 7.**
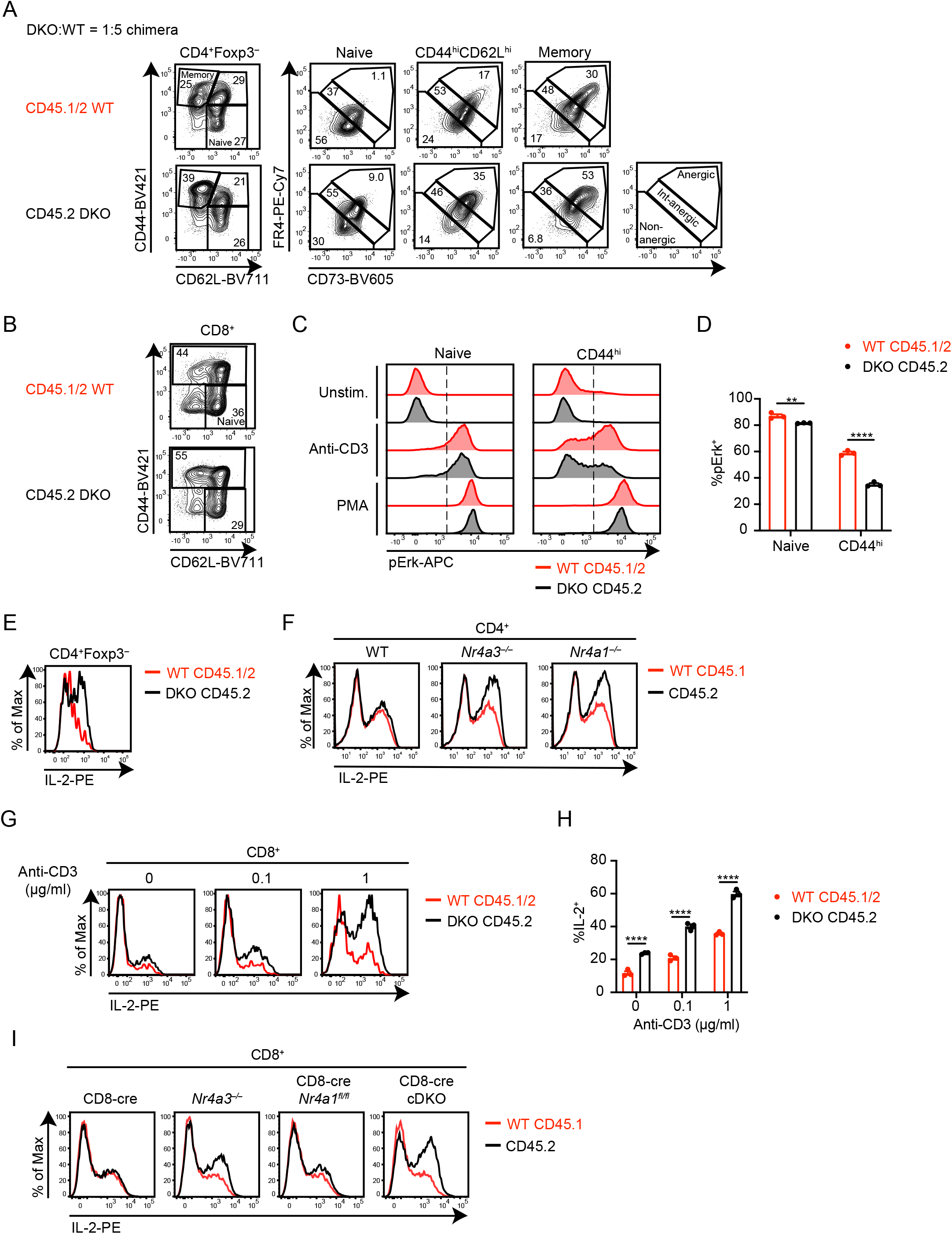
A. Gating strategy corresponding to Fig. 7A to identify splenic CD4+ T cell sub-populations for phosflow assay. Plots are representative of 3 mice. B, C. Splenocytes from DKO:WT = 1:5 chimera were stimulated and stained as described in Fig. 7A. (B) Representative plots show naïve and CD44^hi^CD8^+^ T cell gates. (C) Representative histograms show intracellular pErk expression in naïve and CD44^hi^CD8^+^ T cells. Vertical dashed line shows the threshold of positive gate. Plots are representative of 6 mice. D. Quantification of %pErk^+^ as described in C above (n = 3 biological replicates, representative of n = 2 independent experiments). E. Lymph node cells from DKO:WT = 1:1 chimera were cultured with 0.1 μg/ml of plate-bound anti-CD3 and treated as described for Fig. 7C. Cell surface was stained with CD4, CD8, CD45.1 and CD45.2, followed by permeabilization and intracellular staining for FOXP3 and IL-2. Shown is a representative histogram pre-gated on CD4^+^FOXP3^−^ of each genotype. F. Lymph node cells from WT, *Nr4a3^−/−^* and *Nr4a1^−/−^* mice were mixed with CD45.1 lymph node cells, then cultured with 0.1 μg/ml of plate-bound anti-CD3 and stained as described for Fig. 7C. Plots are representative of 3 mice. G, H. Lymph node cells from 10 weeks post-transplant DKO:WT = 1:1 chimera were stimulated and stained as described in Fig. 7C. (G) Representative histograms show intracellular IL-2 in CD8^+^ T cells of each genotype. (H) Quantification of %IL-2^+^ in CD8^+^ T cells (data in G, H represent n = 3 biological replicates). I. Lymph node cells from CD8-cre, *Nr4a3^−/−^*, CD8-cre *Nr4a1^fl/fl^* or CD8-cre *Nr4a1^fl/fl^ Nr4a3^−/−^* (cDKO) were mixed with CD45.1 lymph node cells, cultured with 1 μg/ml of plate-bound anti-CD3, and then treated and stained as described for Fig. 7C. Representative histograms show intracellular IL-2 in CD8^+^ T cells of each genotype. Plots are representative of 2 independent experiments. Graphs depict mean +/- SEM. Statistical significance was assessed by two-tailed unpaired Student’s t-test with the Holm–Šídák method (D, H). **P < 0.01; ***P < 0.001; ****P < 0.0001.

**Supplementary Figure 8. Supporting data for Figure 8.**
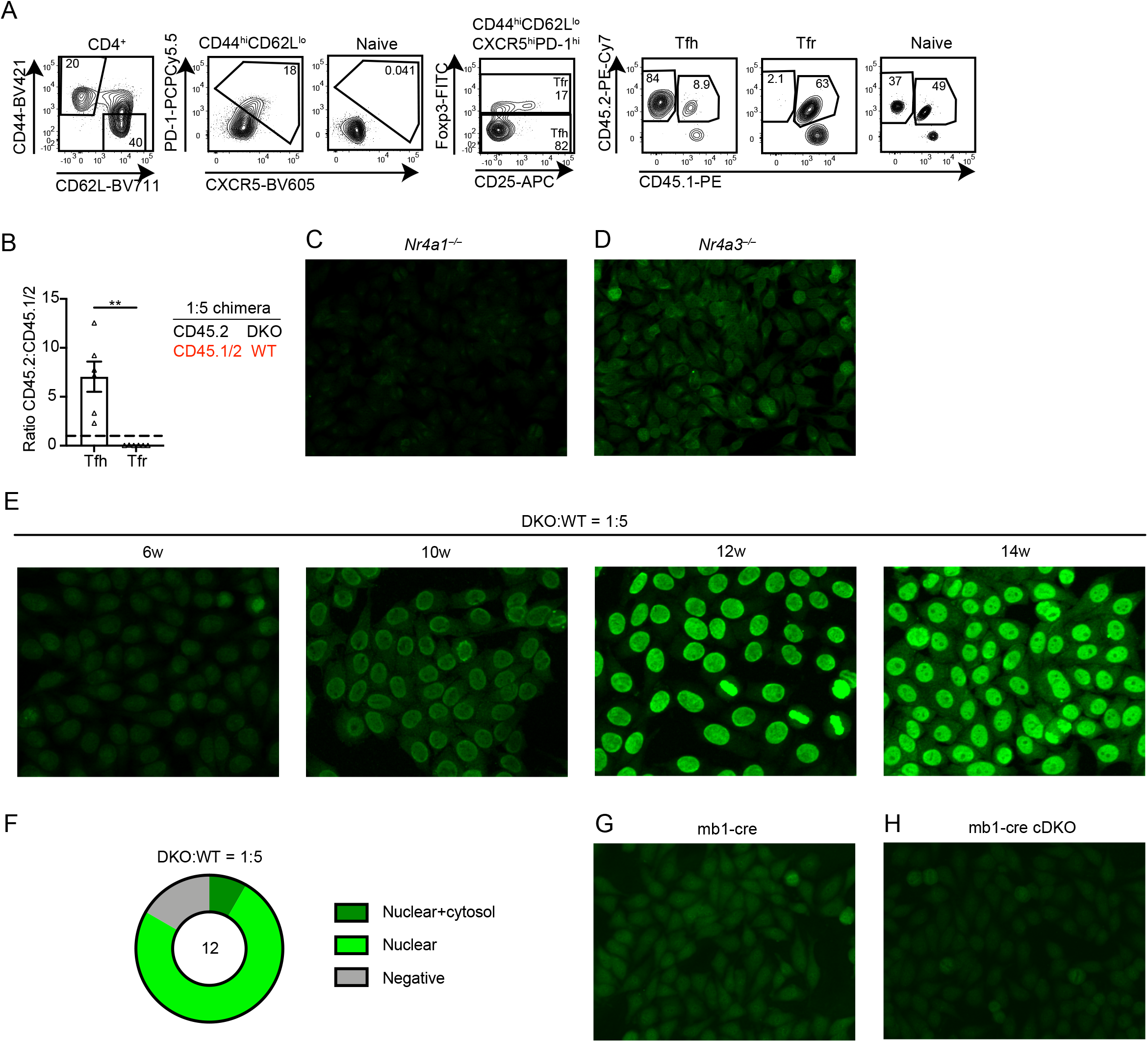
A. Splenocytes from DKO:WT = 1:5 chimera were harvested and stained with CD4, CD25, CD44, CD62L, CXCR5 and PD-1, then permeabilized and stained with FOXP3. Representative plots show FOXP3^−^ follicular helper T cells (Tfh) and FOXP3^+^ follicular regulatory T cells (Tfr) within CD4^+^CD44^hi^CD62L^lo^CXCR5^hi^PD-1^hi^ gate. Naïve CD4^+^ T cell gate is shown for reference to define Tfh/Tfr gate. Donor genotype gates among each population are shown. B. Ratio of CD45.2 to CD45.1/2 for Tfh or Tfr, normalized to naïve CD4^+^ as gated in A above (n = 3 biological replicates). Statistical significance was assessed by two-tailed unpaired Student’s t-test. **P < 0.01. C, D, G, H. Anti-nuclear antibody (ANA) immunofluorescence images as described for Fig. 8A-D. Serum was collected from 12-week-old *Nr4a1^−/−^* (C), *Nr4a3^−/−^* (D) mice, from DKO:WT = 1:5 chimera at 6, 10, 12, or 14 weeks post-transplant, and from mb1-cre *Nr4a1^fl/fl^ Nr4a3^−/−^* (mb1cre-cDKO) (G) or mb1-cre (H) non-competitive chimeras 40 weeks post-transplant. Images are representative of n = 3 (mb1-cre chimera and DKO:WT = 1:5 chimera), n = 5 (mb1-cre cDKO chimera) or n = 10 (*Nr4a1^−/−^* and *Nr4a3^−/−^*) biological replicates. F. Graph depict frequency of Hep-2 staining patterns in DKO:WT = 1:5 chimera.

**Supplementary Figure 9.**
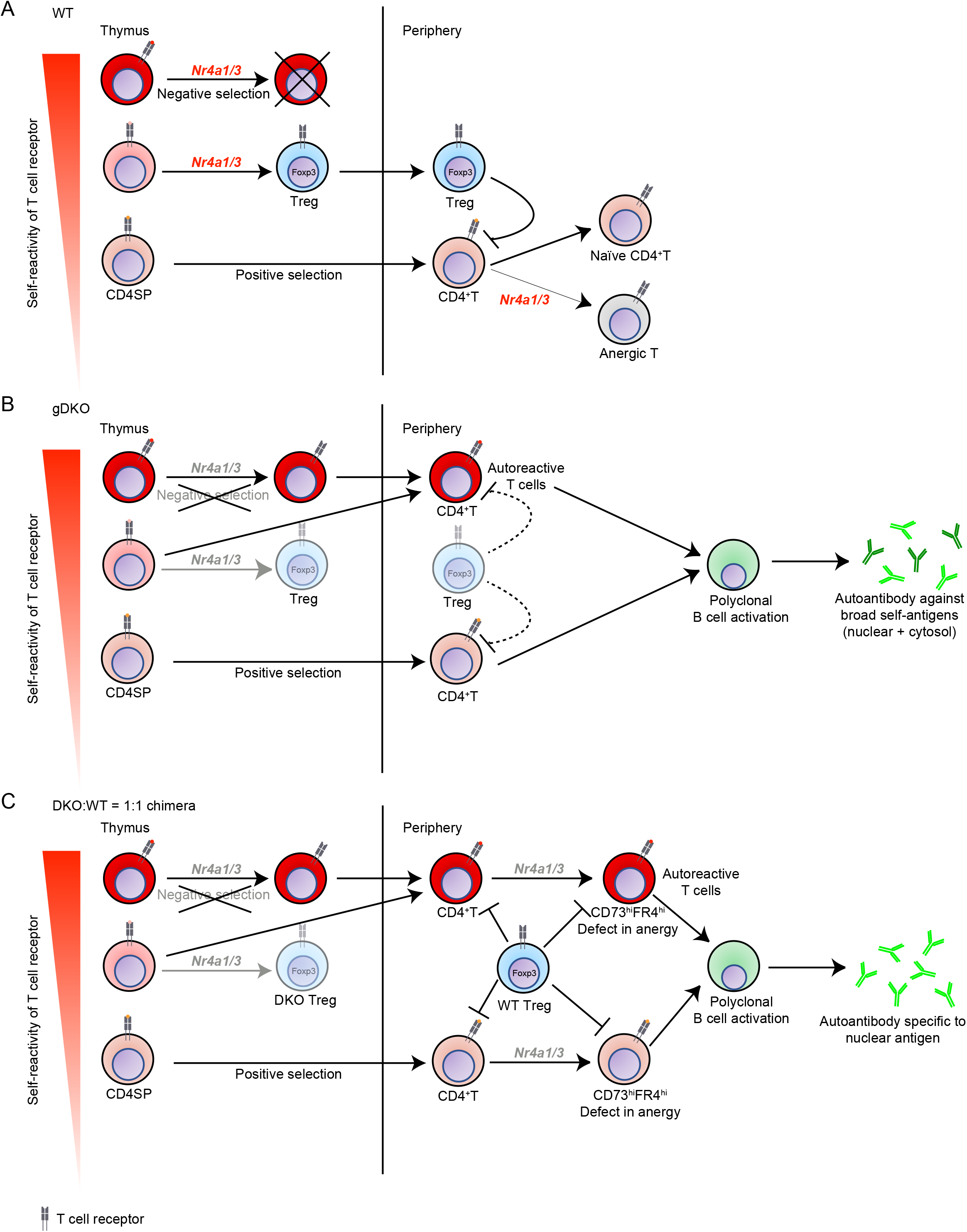
Model. Roles for NR4A family in T cell tolerance and immune homeostasis. A. In WT, central and peripheral tolerance is intact. *Nr4a1* and *Nr4a3* mediate thymic negative selection and Treg homeostasis. *Nr4a1* and *Nr4a3* also play a role on induction and maintenance of anergy in periphery. B. In *Nr4a1^−/−^ Nr4a3^−/−^* (gDKO) mice, both thymic negative selection and Treg homeostasis are impaired, and highly autoreactive T cells escape to periphery. Autoreactive CD4^+^ T cells are activated in part because of Treg deficiency. Polyclonal B cells are activated by autoreactive T cells and produce autoantibody against a broad spectrum of self-antigens as detected by both cytoplasmic and nuclear Hep-2 staining pattern. C. Treg compartment is reconstituted by WT donor BM in DKO:WT competitive chimeras. However, negative selection of self-reactive DKO thymocytes is still impaired. Self-reactive DKO T cells that escaped negative selection encounter self-antigens in the periphery, acquire features of anergy including expression of CD73 and FR4, yet exhibit a defect in peripheral tolerance. WT-origin Treg and cell-intrinsic peripheral tolerance mechanisms are insufficient to completely suppress autoreactive DKO T cells, which in turn drive anti-nuclear autoantibody production. Although Treg compartment restores some immune homeostasis (and tolerance to cytosolic antigens as detected by Hep-2 staining), autoimmunity is not suppressed due to role of NR4A family in other T cell-intrinsic tolerance mechanisms.

## Notes

### Competing Interest Statement

JZ serves on SAB for Walking Fish Therapeutics

## References

1. Sakaguchi S, Sakaguchi N, Asano M, Itoh M, and Toda M. Immunologic self-tolerance maintained by activated T cells expressing IL-2 receptor alpha-chains (CD25). Breakdown of a single mechanism of self-tolerance causes various autoimmune diseases. J Immunol. 1995;155(3):1151–64.

2. Sakaguchi S. Naturally arising Foxp3-expressing CD25+CD4+ regulatory T cells in immunological tolerance to self and non-self. Nat Immunol. 2005;6(4):345–52.

3. Gavin MA, Rasmussen JP, Fontenot JD, Vasta V, Manganiello VC, Beavo JA, et al. Foxp3-dependent programme of regulatory T-cell differentiation. Nature. 2007;445(7129):771–5.

4. Lahl K, Loddenkemper C, Drouin C, Freyer J, Arnason J, Eberl G, et al. Selective depletion of Foxp3+ regulatory T cells induces a scurfy-like disease. J Exp Med. 2007;204(1):57–63.

5. Kim JM, Rasmussen JP, and Rudensky AY. Regulatory T cells prevent catastrophic autoimmunity throughout the lifespan of mice. Nat Immunol. 2007;8(2):191–7.

6. Fontenot JD, Gavin MA, and Rudensky AY. Foxp3 programs the development and function of CD4+CD25+ regulatory T cells. Nat Immunol. 2003;4(4):330–6.

7. Ramsdell F, and Ziegler SF. FOXP3 and scurfy: how it all began. Nat Rev Immunol. 2014;14(5):343–9.

8. Hadaschik EN, Wei X, Leiss H, Heckmann B, Niederreiter B, Steiner G, et al. Regulatory T cell-deficient scurfy mice develop systemic autoimmune features resembling lupus-like disease. Arthritis Res Ther. 2015;17:35.

9. Skuljec J, Cabanski M, Surdziel E, Lachmann N, Brennig S, Pul R, et al. Monocyte/macrophage lineage commitment and distribution are affected by the lack of regulatory T cells in scurfy mice. Eur J Immunol. 2016;46(7):1656–68.

10. Passerini L, Rossi Mel E, Sartirana C, Fousteri G, Bondanza A, Naldini L, et al. CD4(+) T cells from IPEX patients convert into functional and stable regulatory T cells by FOXP3 gene transfer. Science translational medicine. 2013;5(215):215ra174.

11. Cambier JC, Gauld SB, Merrell KT, and Vilen BJ. B-cell anergy: from transgenic models to naturally occurring anergic B cells? Nat Rev Immunol. 2007;7(8):633–43.

12. Nemazee D. Mechanisms of central tolerance for B cells. Nat Rev Immunol. 2017;17(5):281–94.

13. ElTanbouly MA, and Noelle RJ. Rethinking peripheral T cell tolerance: checkpoints across a T cell’s journey. Nat Rev Immunol. 2021;21(4):257–67.

14. Sekiya T, Kashiwagi I, Yoshida R, Fukaya T, Morita R, Kimura A, et al. Nr4a receptors are essential for thymic regulatory T cell development and immune homeostasis. Nat Immunol. 2013;14(3):230–7.

15. Sekiya T, Hibino S, Saeki K, Kanamori M, Takaki S, and Yoshimura A. Nr4a Receptors Regulate Development and Death of Labile Treg Precursors to Prevent Generation of Pathogenic Self-Reactive Cells. Cell Rep. 2018;24(6):1627–38 e6.

16. Ashouri JF, Hsu LY, Yu S, Rychkov D, Chen Y, Cheng DA, et al. Reporters of TCR signaling identify arthritogenic T cells in murine and human autoimmune arthritis. Proc Natl Acad Sci U S A. 2019;116(37):18517–27.

17. Chen J, Lopez-Moyado IF, Seo H, Lio CJ, Hempleman LJ, Sekiya T, et al. NR4A transcription factors limit CAR T cell function in solid tumours. Nature. 2019;567(7749):530–4.

18. Boudreaux SP, Duren RP, Call SG, Nguyen L, Freire PR, Narayanan P, et al. Drug targeting of NR4A nuclear receptors for treatment of acute myeloid leukemia. Leukemia. 2019;33(1):52–63.

19. Winoto A, and Littman DR. Nuclear hormone receptors in T lymphocytes. Cell. 2002;109 Suppl:S57-66.

20. Calnan BJ, Szychowski S, Chan FK, Cado D, and Winoto A. A role for the orphan steroid receptor Nur77 in apoptosis accompanying antigen-induced negative selection. Immunity. 1995;3(3):273–82.

21. Zikherman J, Parameswaran R, and Weiss A. Endogenous antigen tunes the responsiveness of naive B cells but not T cells. Nature. 2012;489(7414):160–4.

22. Tan C, Mueller JL, Noviski M, Huizar J, Lau D, Dubinin A, et al. Nur77 Links Chronic Antigen Stimulation to B Cell Tolerance by Restricting the Survival of Self-Reactive B Cells in the Periphery. J Immunol. 2019;202(10):2907–23.

23. Au-Yeung BB, Melichar HJ, Ross JO, Cheng DA, Zikherman J, Shokat KM, et al. Quantitative and temporal requirements revealed for Zap70 catalytic activity during T cell development. Nat Immunol. 2014;15(7):687–94.

24. Liu X, Wang Y, Lu H, Li J, Yan X, Xiao M, et al. Genome-wide analysis identifies NR4A1 as a key mediator of T cell dysfunction. Nature. 2019;567(7749):525-9.

25. Moran AE, Holzapfel KL, Xing Y, Cunningham NR, Maltzman JS, Punt J, et al. T cell receptor signal strength in Treg and iNKT cell development demonstrated by a novel fluorescent reporter mouse. J Exp Med. 2011;208(6):1279–89.

26. Zinzow-Kramer WM, Weiss A, and Au-Yeung BB. Adaptation by naive CD4(+) T cells to self-antigen-dependent TCR signaling induces functional heterogeneity and tolerance. Proc Natl Acad Sci U S A. 2019;116(30):15160–9.

27. Tan C, Hiwa R, Mueller JL, Vykunta V, Hibiya K, Noviski M, et al. NR4A nuclear receptors restrain B cell responses to antigen when second signals are absent or limiting. Nat Immunol. 2020;21(10):1267–79.

28. Zhou T, Cheng J, Yang P, Wang Z, Liu C, Su X, et al. Inhibition of Nur77/Nurr1 leads to inefficient clonal deletion of self-reactive T cells. The Journal of experimental medicine. 1996;183(4):1879–92.

29. Cheng LE, Chan FK, Cado D, and Winoto A. Functional redundancy of the Nur77 and Nor-1 orphan steroid receptors in T-cell apoptosis. The EMBO journal. 1997;16(8):1865–75.

30. Lee SL, Wesselschmidt RL, Linette GP, Kanagawa O, Russell JH, and Milbrandt J. Unimpaired thymic and peripheral T cell death in mice lacking the nuclear receptor NGFI-B (Nur77). *Science (New York*, NY). 1995;269(5223):532–5.

31. Fassett MS, Jiang W, D’Alise AM, Mathis D, and Benoist C. Nuclear receptor Nr4a1 modulates both regulatory T-cell (Treg) differentiation and clonal deletion. Proc Natl Acad Sci U S A. 2012;109(10):3891–6.

32. Mullican SE, Zhang S, Konopleva M, Ruvolo V, Andreeff M, Milbrandt J, et al. Abrogation of nuclear receptors Nr4a3 and Nr4a1 leads to development of acute myeloid leukemia. Nat Med. 2007;13(6):730–5.

33. Koenis DS, Medzikovic L, Vos M, Beldman TJ, van Loenen PB, van Tiel CM, et al. Nur77 variants solely comprising the amino-terminal domain activate hypoxia-inducible factor-1alpha and affect bone marrow homeostasis in mice and humans. J Biol Chem. 2018;293(39):15070–83.

34. Lio CW, and Hsieh CS. A two-step process for thymic regulatory T cell development. Immunity. 2008;28(1):100–11.

35. Palacios EH, and Weiss A. Distinct roles for Syk and ZAP-70 during early thymocyte development. J Exp Med. 2007;204(7):1703–15.

36. Taghon T, Yui MA, Pant R, Diamond RA, and Rothenberg EV. Developmental and molecular characterization of emerging beta- and gammadelta-selected pre- T cells in the adult mouse thymus. Immunity. 2006;24(1):53–64.

37. Baldwin TA, and Hogquist KA. Transcriptional analysis of clonal deletion in vivo. J Immunol. 2007;179(2):837–44.

38. Liu ZG, Smith SW, McLaughlin KA, Schwartz LM, and Osborne BA. Apoptotic signals delivered through the T-cell receptor of a T-cell hybrid require the immediate-early gene nur77. Nature. 1994;367(6460):281–4.

39. Woronicz JD, Calnan B, Ngo V, and Winoto A. Requirement for the orphan steroid receptor Nur77 in apoptosis of T-cell hybridomas. Nature. 1994;367(6460):277–81.

40. Nowyhed HN, Huynh TR, Blatchley A, Wu R, Thomas GD, and Hedrick CC. The nuclear receptor nr4a1 controls CD8 T cell development through transcriptional suppression of runx3. Sci Rep. 2015;5:9059.

41. Ramirez-Herrick AM, Mullican SE, Sheehan AM, and Conneely OM. Reduced NR4A gene dosage leads to mixed myelodysplastic/myeloproliferative neoplasms in mice. Blood. 2011;117(9):2681–90.

42. Clark LB, Appleby MW, Brunkow ME, Wilkinson JE, Ziegler SF, and Ramsdell F. Cellular and molecular characterization of the scurfy mouse mutant. Journal of immunology (Baltimore, Md : 1950). 1999;162(5):2546-54.

43. Fontenot JD, Rasmussen JP, Williams LM, Dooley JL, Farr AG, and Rudensky AY. Regulatory T cell lineage specification by the forkhead transcription factor foxp3. Immunity. 2005;22(3):329–41.

44. Chinen T, Kannan AK, Levine AG, Fan X, Klein U, Zheng Y, et al. An essential role for the IL-2 receptor in Treg cell function. Nat Immunol. 2016;17(11):1322–33.

45. Spolski R, Li P, and Leonard WJ. Biology and regulation of IL-2: from molecular mechanisms to human therapy. Nat Rev Immunol. 2018;18(10):648–59.

46. Maekawa Y, Minato Y, Ishifune C, Kurihara T, Kitamura A, Kojima H, et al. Notch2 integrates signaling by the transcription factors RBP-J and CREB1 to promote T cell cytotoxicity. Nat Immunol. 2008;9(10):1140–7.

47. Martinez RJ, Zhang N, Thomas SR, Nandiwada SL, Jenkins MK, Binstadt BA, et al. Arthritogenic self-reactive CD4+ T cells acquire an FR4hiCD73hi anergic state in the presence of Foxp3+ regulatory T cells. J Immunol. 2012;188(1):170–81.

48. Kalekar LA, Schmiel SE, Nandiwada SL, Lam WY, Barsness LO, Zhang N, et al. CD4(+) T cell anergy prevents autoimmunity and generates regulatory T cell precursors. Nat Immunol. 2016;17(3):304–14.

49. Fathman CG, and Lineberry NB. Molecular mechanisms of CD4+ T-cell anergy. Nat Rev Immunol. 2007;7(8):599–609.

50. Zheng Y, Zha Y, and Gajewski TF. Molecular regulation of T-cell anergy. EMBO Rep. 2008;9(1):50–5.

51. Wells AD. New insights into the molecular basis of T cell anergy: anergy factors, avoidance sensors, and epigenetic imprinting. J Immunol. 2009;182(12):7331–41.

52. Aschermann S, Lehmann CH, Mihai S, Schett G, Dudziak D, and Nimmerjahn F. B cells are critical for autoimmune pathology in Scurfy mice. Proc Natl Acad Sci U S A. 2013;110(47):19042–7.

53. Haeberle S, Raker V, Haub J, Kim YO, Weng SY, Yilmaz OK, et al. Regulatory T cell deficient scurfy mice exhibit a Th2/M2-like inflammatory response in the skin. J Dermatol Sci. 2017;87(3):285–91.

54. Yilmaz OK, Haeberle S, Zhang M, Fritzler MJ, Enk AH, and Hadaschik EN. Scurfy Mice Develop Features of Connective Tissue Disease Overlap Syndrome and Mixed Connective Tissue Disease in the Absence of Regulatory T Cells. Front Immunol. 2019;10:881.

55. Vinuesa CG, Linterman MA, Yu D, and MacLennan IC. Follicular Helper T Cells. Annu Rev Immunol. 2016;34:335–68.

56. Stebegg M, Kumar SD, Silva-Cayetano A, Fonseca VR, Linterman MA, and Graca L. Regulation of the Germinal Center Response. Front Immunol. 2018;9(2469):2469.

57. Ma W, Zhao R, Yang R, Liu B, Chen X, Wu L, et al. Nuclear receptors of the NR4a family are not required for the development and function of follicular T helper cells. Int Immunopharmacol. 2015;28(2):841–5.

58. Sekiya T, Kondo T, Shichita T, Morita R, Ichinose H, and Yoshimura A. Suppression of Th2 and Tfh immune reactions by Nr4a receptors in mature T reg cells. J Exp Med. 2015;212(10):1623–40.

59. Sekiya T, Kagawa S, Masaki K, Fukunaga K, Yoshimura A, and Takaki S. Regulation of peripheral Th/Treg differentiation and suppression of airway inflammation by Nr4a transcription factors. iScience. 2021;24(3):102166.

60. Hu QN, and Baldwin TA. Differential roles for Bim and Nur77 in thymocyte clonal deletion induced by ubiquitous self-antigen. J Immunol. 2015;194(6):2643–53.

61. Stritesky GL, Xing Y, Erickson JR, Kalekar LA, Wang X, Mueller DL, et al. Murine thymic selection quantified using a unique method to capture deleted T cells. Proc Natl Acad Sci U S A. 2013;110(12):4679–84.

62. Klein L, Robey EA, and Hsieh CS. Central CD4(+) T cell tolerance: deletion versus regulatory T cell differentiation. Nat Rev Immunol. 2019;19(1):7–18.

63. Suen AY, and Baldwin TA. Proapoptotic protein Bim is differentially required during thymic clonal deletion to ubiquitous versus tissue-restricted antigens. Proc Natl Acad Sci U S A. 2012;109(3):893–8.

64. Thompson J, and Winoto A. During negative selection, Nur77 family proteins translocate to mitochondria where they associate with Bcl-2 and expose its proapoptotic BH3 domain. J Exp Med. 2008;205(5):1029–36.

65. Odagiu L, May J, Boulet S, Baldwin TA, and Labrecque N. Role of the Orphan Nuclear Receptor NR4A Family in T-Cell Biology. Front Endocrinol (Lausanne). 2020;11:624122.

66. Alam A, Braun MY, Hartgers F, Lesage S, Cohen L, Hugo P, et al. Specific activation of the cysteine protease CPP32 during the negative selection of T cells in the thymus. J Exp Med. 1997;186(9):1503–12.

67. Bouillet P, Purton JF, Godfrey DI, Zhang LC, Coultas L, Puthalakath H, et al. BH3-only Bcl-2 family member Bim is required for apoptosis of autoreactive thymocytes. Nature. 2002;415(6874):922–6.

68. Lin B, Kolluri SK, Lin F, Liu W, Han YH, Cao X, et al. Conversion of Bcl-2 from protector to killer by interaction with nuclear orphan receptor Nur77/TR3. Cell. 2004;116(4):527–40.

69. Zarraga-Granados G, Mucino-Hernandez G, Sanchez-Carbente MR, Villamizar-Galvez W, Penas-Rincon A, Arredondo C, et al. The nuclear receptor NR4A1 is regulated by SUMO modification to induce autophagic cell death. PLoS One. 2020;15(3):e0222072.

70. Odagiu L, Boulet S, Maurice De Sousa D, Daudelin JF, Nicolas S, and Labrecque N. Early programming of CD8(+) T cell response by the orphan nuclear receptor NR4A3. Proc Natl Acad Sci U S A. 2020;117(39):24392–402.

71. McEvoy C, de Gaetano M, Giffney HE, Bahar B, Cummins EP, Brennan EP, et al. NR4A Receptors Differentially Regulate NF-kappaB Signaling in Myeloid Cells. Front Immunol. 2017;8:7.

72. Zhou H, Wang J, Zhu P, Zhu H, Toan S, Hu S, et al. NR4A1 aggravates the cardiac microvascular ischemia reperfusion injury through suppressing FUNDC1- mediated mitophagy and promoting Mff-required mitochondrial fission by CK2alpha. Basic Res Cardiol. 2018;113(4):23.

73. Wang LM, Zhang Y, Li X, Zhang ML, Zhu L, Zhang GX, et al. Nr4a1 plays a crucial modulatory role in Th1/Th17 cell responses and CNS autoimmunity. Brain Behav Immun. 2018;68:44–55.

74. Liebmann M, Hucke S, Koch K, Eschborn M, Ghelman J, Chasan AI, et al. Nur77 serves as a molecular brake of the metabolic switch during T cell activation to restrict autoimmunity. Proc Natl Acad Sci U S A. 2018;115(34):E8017–E26.

75. Zhan Y, Du X, Chen H, Liu J, Zhao B, Huang D, et al. Cytosporone B is an agonist for nuclear orphan receptor Nur77. Nat Chem Biol. 2008;4(9):548–56.

76. Zhan YY, Chen Y, Zhang Q, Zhuang JJ, Tian M, Chen HZ, et al. The orphan nuclear receptor Nur77 regulates LKB1 localization and activates AMPK. Nat Chem Biol. 2012;8(11):897–904.

77. Hobeika E, Thiemann S, Storch B, Jumaa H, Nielsen PJ, Pelanda R, et al. Testing gene function early in the B cell lineage in mb1-cre mice. Proc Natl Acad Sci U S A. 2006;103(37):13789–94.

